# A Volatile Signal Controls Virulence in the Plant Pathogen *Pseudomonas syringae* pv. *syringae* and a Strategy for Infection Control in Organic Farming

**DOI:** 10.1101/2020.09.18.279364

**Authors:** Simon Sieber, Anugraha Mathew, Christian Jenul, Tobias Kohler, Max Bär, Víctor J. Carrión, Francisco M. Cazorla, Urs Stalder, Ya-Chu Hsieh, Laurent Bigler, Leo Eberl, Karl Gademann

## Abstract

*Pseudomonas syringae* is an important pathogen of many agriculturally valuable crops. Among the various pathovars described *P. syringae* pv. *syringae* (Pss) has a particularly wide host range, infecting primarily woody and herbaceous host plants. The ability of Pss to cause bacterial apical necrosis of mango trees is dependent on the production of the antimetabolite toxin mangotoxin. The production of this toxin was shown to be regulated by a self-produced signaling molecule. In this study, we determined the structure of the Pss signal molecule belonging to the recently described family of diazeniumdiolate communication molecules. Employing a targeted mass spectrometry-based approach, we provide experimental evidence that the major signal produced by Pss is the volatile compound leudiazen, which controls mangotoxin production and virulence in a detached tomato leaflet infection model. Experimental results demonstrate that KMnO_4_ solution inactivates leudiazen and that treatment of infected leaves with KMnO_4_ abolishes necrosis. This strategy represents the first example of chemically degrading a signaling molecule to interfere with bacterial communication. The application of KMnO_4_ solution, which is regulatorily approved in organic farming, may constitute an environmentally friendly strategy to control Pss infections.

## Introduction

Bacteria have the extraordinary ability to sense and respond to their population density. This is generally achieved by excreting small molecules that act as messengers. When a critical threshold concentration of the signal molecule is reached, it interacts with a cognate receptor, resulting in the activation or repression of target genes.^1^ Adaptation and coordination of gene expression is particularly important for pathogenic microorganisms that need to overcome the innate immune system of their hosts.

Among the Gram-negative bacteria the most commonly used signals for monitoring their own population density are *N*-acyl-homoserine lactones (AHLs), a phenomenon that has been termed quorum sensing.^2^ AHL-dependent quorum sensing (QS) systems have been identified in more than 200 species, in which they regulate a wide variety of functions, often in connection with pathogenicity, surface colonization, and antibiotic production.^3^

Representatives of the genus *Pseudomonas* represent a good example of both benign and pathogenic plant–bacteria interactions.^4^ While some strains are beneficial for plants,^5^ many others, such as members of the genus *Pseudomonas syringae* (*P. syringae*), are important pathogens, which infect almost all economically important crop species.^6^ Decades of studies have revealed the broad repertoire of virulence strategies employed by *P. syringae*, which include large numbers of functionally redundant type III secretion system (T3SS) effectors and phytotoxins.^7^ While the major lipodepsipeptide toxins, such as the syringomycins and syringopeptins, cause direct damage to plant cells,^8,9^ the modified peptide toxins, phaseolotoxin^10,11^ and tabtoxin,^12^ specifically target enzymes involved in amino acid biosynthesis.^13^

Another toxin that has an important ecological impact is mangotoxin, which is an inhibitor of ornithine biosynthesis.^14^ This compound has been isolated from *P. syringae* pv. *syringae* (Pss) which is the causal agent of bacterial apical necrosis (BAN) of mango trees.^15^ To date, the structure of mangotoxin is unknown but a preliminary characterization indicated that the molecular weight of mangotoxin could be smaller than 3 kDa and that the compound could be resistant to extreme pH and high temperature.^14^

The *mbo* gene cluster encoding the mangotoxin biosynthesis genes was first described in Pss^16^ and was later also identified in many other *P. syringae* pathovars.^17–19^ Investigations of the regulation of mangotoxin production has led to the identification of the *mgo* operon, which was shown to be required for the production of an unknown signal molecule.^20,21^ The *mgo* gene cluster comprises four genes, namely *mgoB*, a predicted haem oxygenase, *mgoC*, a homolog of the *N*-oxygenase *aurF*,^22^ *mgoA*, a non-ribosomal peptide synthetase (NRPS) and *mgoD* a putative polyketide cyclase/dehydratase.^20,21^ Homologs of the *mgo* cluster have also been identified in other *Pseudomonas* strains such as the insect pathogen *Pseudomonas entomophila* (*pvf* cluster),^23^ the sugar cane endophyte *Pseudomonas aurantiaca* PB-St2,^24^ *Pseudomonas* sp. SH-C52,^25^ several *Pseudomonas fluorescens* strains,^26^ and from various pathogenic *Burkholderia* species (*ham* cluster).^27,28^

Recently, we discovered that the *ham* gene cluster directs the synthesis of two diazeniumdiolate natural products: the antifungal metabolite fragin and the novel signaling molecule valdiazen (**1, Figure 1**).^27^ During the last two years, this functional group has been also associated with the discovery of a new class of siderophore^29–31^ and the identification of alanosine gene cluster.^32,33^ In *Burkholderia cenocepacia* H111, *hamA, C, D*, and *E* genes, which are homologs of the *mgo* genes, were shown to be involved in the biosynthesis of valdiazen (**1**)^27,34^ The proposed biosynthesis of valdiazen (**1**) starts with L-valine as the initial NRPS substrate and the diazeniumdiolate functional group is hypothesized to be formed by tailoring enzymes. To conclude, the NRPS reduction domain of HamD is thought to perform the last steps of valdiazen (**1**) biosynthesis with a four electron reduction process.^27,34^ Furthermore, in 2018, Li and coworkers engineered a *P. entomophila* strain to overexpress the *pvf* cluster leading to the production of three *N*-oxide metabolites^35^ and they investigated the role of the *N*-oxygenase in the biosynthesis of valdiazen (**1**).^36^ In a follow up study, the same research group investigated the substrate specificity of the NRPS domain of PvfC and its homologs MgoA and HamD in different proteobacteria and concluded that at least two types of signaling molecules are produced by the different strains^37^

**Figure 1.**
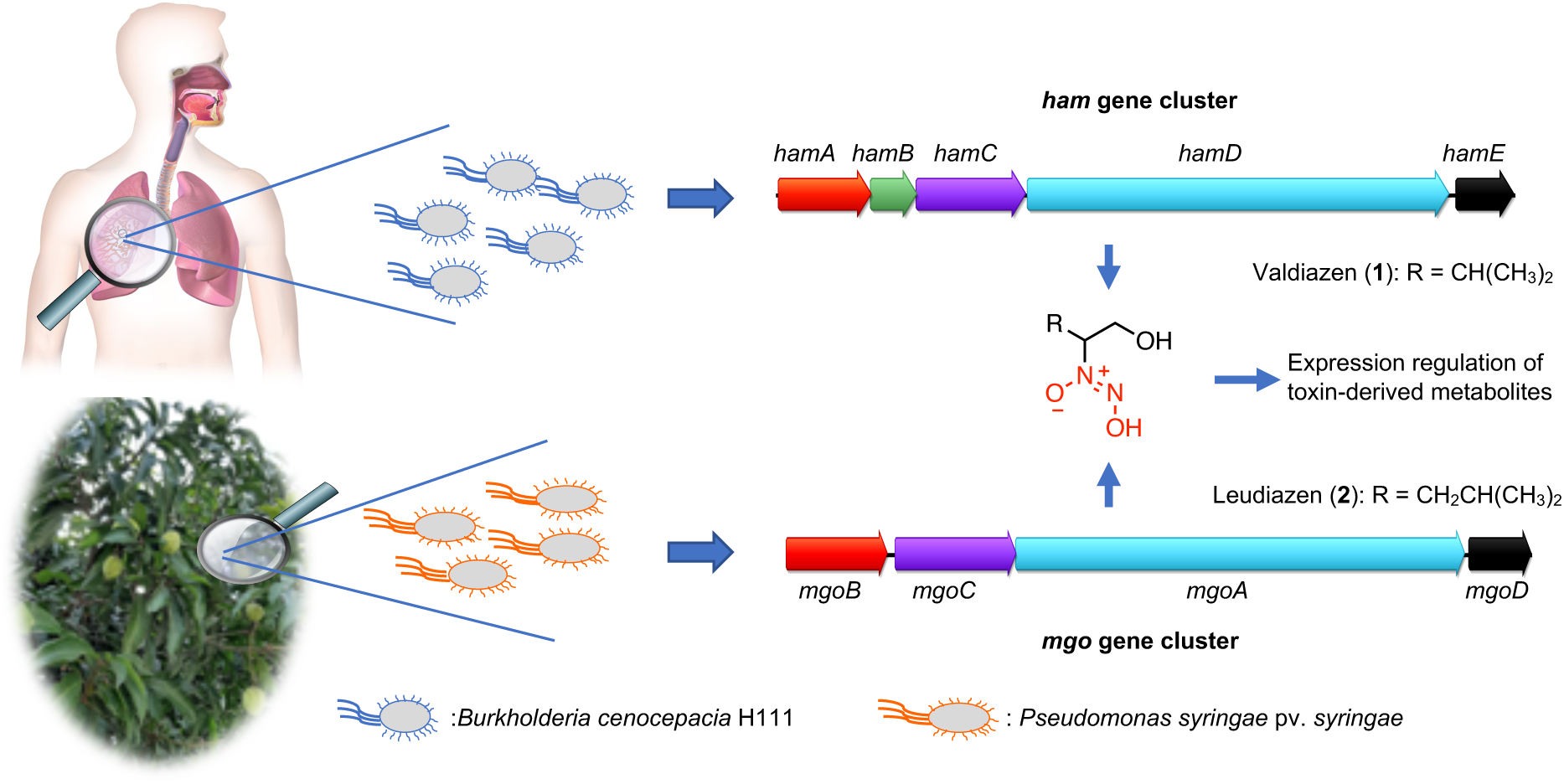
The *ham* gene cluster was identified in the genome of the human pathogen *B. cenocepacia* H111 and the genes *hamA, hamC, hamD* and *hamE* direct the synthesis of diazeniumdiolate valdiazen (**1**).^27,34^ Valdiazen (**1**) regulates the expression of more than 100 genes including the *hamABCDE* and *hamFG* operons encoding the major antifungal metabolite of the strain, fragin. Likewise, the homologous *mgo* cluster in the plant pathogenic bacteria Pss^16,20^ was shown to encode a signaling molecule that regulates the production of mangotoxin.^21^

To date, the signaling molecules produced by *P. entomophila* and Pss have not been identified. Based on our recent discovery of valdiazen (**1**), we aimed at identifying the structure of the signaling molecule produced by Pss, which we hypothesized to be similar to **1**. Furthermore, we speculated that the knowledge of the signaling molecule structure might pave the way to develop a strategy for plant protection (**Figure 1**).

## Results

### Isolation of the Putative Mangotoxin Signaling Compound

Mangotoxin biosynthesis is regulated by a yet unknown signal encoded by the *mgoBCAD* operon.^21^ Accordingly, the activity of the *mboA* promoter was found to be significantly reduced in a *ΔmgoA* mutant background compared to the activity in the wild type strain.^16^ We used a transcriptional fusion of the *mbo* promoter to the *lacZ* gene in the *ΔmgoA* mutant background as a sensor strain for the identification of the unknown signal. Due to the high similarities between the *mgo* gene cluster^20^ and the recently elucidated *ham* gene cluster encoding valdiazen (**1**),^27^ we hypothesized that the signaling molecule produced by Pss may be valdiazen (**1**) or a related compound. In order to isolate the signal compound, we employed a similar isolation strategy as for valdiazen (**1**).^27^ This included an acidification of the supernatant to a pH of 5 followed by a liquid-liquid extraction with CH_2_Cl_2_. The isogenic *ΔmgoA* mutant (*ΔmgoA*)^20^ was used as negative control. We confirmed that the mangotoxin promoter was induced by the extract of the wild type but not by the *ΔmgoA* (**Figure 2A**). Valdiazen (**1**) was found to only weakly induce the biosensor relative to the Pss extract, indicating that the unknown signaling molecule is likely not valdiazen (**1**) but a structurally related molecule (**Figure 2A**).

**Figure 2.**
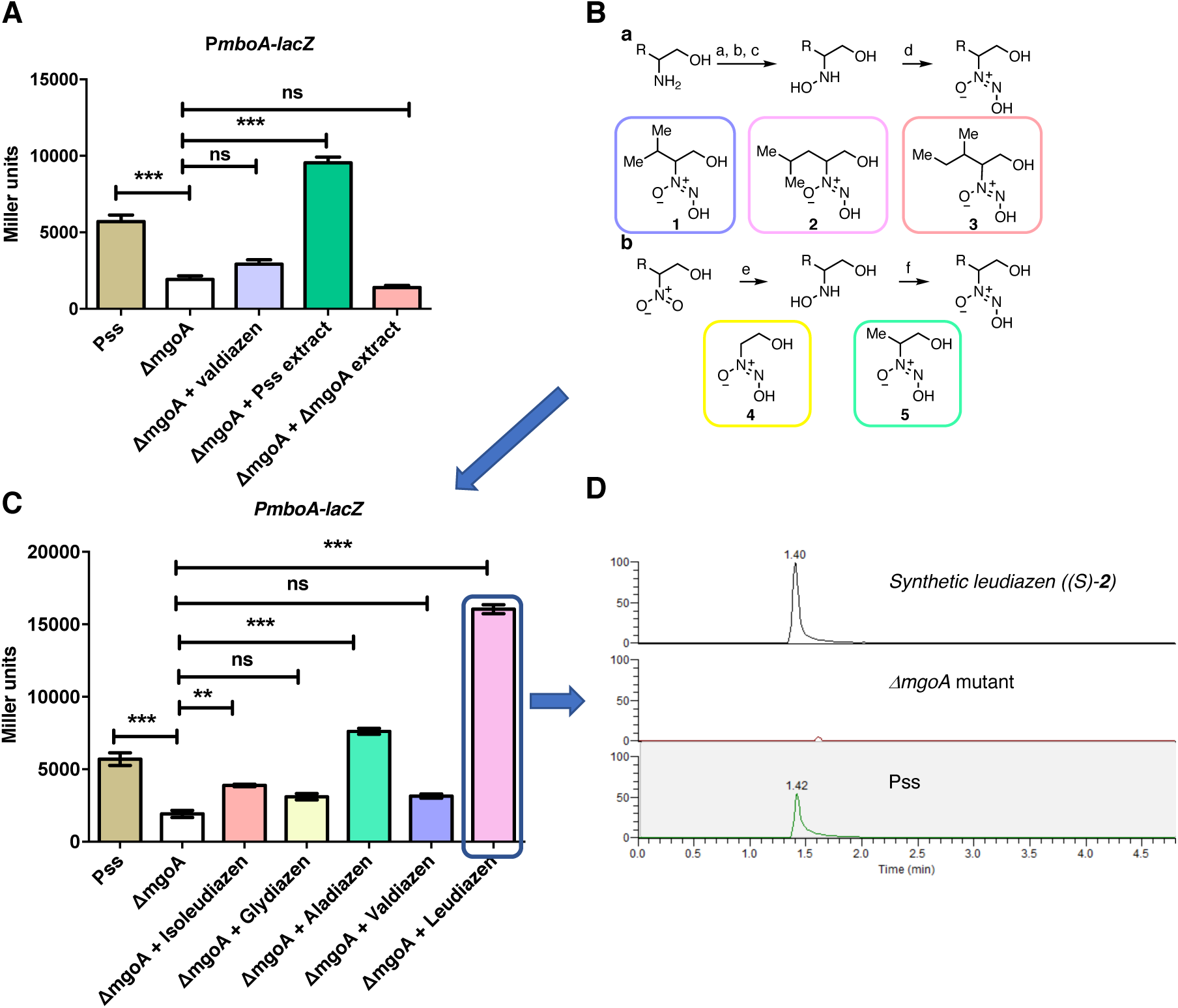
**A**) The *mbo* promoter is activated by a compound present in the extract of the Pss supernatant. The activity of the P*mboA*-*lacZ* fusion was measured in the presence of extracts of the wild type (Pss) and the *ΔmgoA* mutant strain as well as in the presence of synthetic valdiazen (**1**, 50 μM). As a control, we also included the wild type strain (Pss) harboring the sensor plasmid. Results are presented as mean and error bars represent SE. Statistical analysis was performed using one-way ANOVA and Tukey’s multiple comparison as post-test; ns, not significant. Significance is indicated by two or three stars. \*\*\**p* < 0.001, *n* = 3; **B**) **a**) Synthetic route for leudiazen (**2**) and isoleudiazen (**3**). a) *p*-anisaldehyde, CH_2_Cl_2_, RT, 45 min, 40°C, 1.5 h; b) *m*-CPBA, CH_2_Cl_2,_ 0°C, 1 h, RT, 1 h; H_2_NOH.HCl, MeOH, RT, 3 d; d) isoamyl nitrite, MeOH, NH_3,(g)_, RT, 30 min, 15% over 4 steps for leudiazen (**2**) and 10% over 4 steps for isoleudiazen (**3**). (**b**) Synthetic pathway for glydiazen (**4**) and aladiazen (**5**). e) Pd/C (10%), EtOH, RT, 2 h; f) Isoamyl nitrite, 0°C, 15 min, RT, 20 min, 18% over 2 steps for glydiazen (**4**) and 31% over 2 steps for aladiazen (**5**); **C**) The activity of the *PmboA-lacZ* fusion in the presence of different valdiazen derivatives: The β-galactosidase activity of the biosensor plasmid was low in the *ΔmgoA* mutant relative to the activity in the wild type background (Pss) but could be stimulated to different degrees in the presence of valdiazen derivatives. Results are presented as mean and error bars represent SE. Statistical analysis was performed using one-way ANOVA and Tukey’s multiple comparison as post-test; ns, not significant. Significance is indicated by two or three stars. \*\*\**p* < 0.001, *n* = 3; **D**) Specific detection of leudiazen (**2**) with UHPLC-MS/MS in SRM mode (*m/z* 163 to 83 Da, rt = 1.4 min). From top to bottom: synthetic leudiazen ((*S)*-**2**), supernatant of the *ΔmgoA* mutant and supernatant of Pss.

Extracts of the wild type and the *ΔmgoA* mutant were compared using several analytical techniques to identify the signal molecule. Both samples were extensively studied by NMR and UHPLC/MS but no difference could be observed. We hypothesized that the concentration of the signaling molecule was significantly lower than in the case of valdiazen (22 μM)^27^ and we therefore employed several methods to concentrate and purify the compound. A preliminary separation of the extract was achieved using a reverse phase (RP) solid phase extraction (SPE), indicating that the compound was retained in the column with H_2_O as solvent but was eluted with MeOH. Encouraged by the capacity of a reverse phase resin to separate the compound, a fractionation by HPLC was performed using H_2_O and MeCN possessing either 0.1% HCOOH or 0.1% NH_4_OAc as additive. Unfortunately, the obtained fractions possessed a lower activity in our assay than the original extract. These results suggested that the compound was either unstable or volatile.

### Synthesis of Valdiazen Derivatives

Since our results indicated that the compound is potentially an analog of valdiazen (**1**), we embarked on the synthesis of several derivatives of **1**. We selected the aliphatic amino acids glycine, alanine, leucine, and isoleucine as starting materials. The synthesis pathway of leudiazen (**2**) and isoleudiazen (**3**) followed our previously described route.^27^ Briefly, each amino alcohol was oxidized using a three step procedure to obtain the corresponding hydroxylamine.^38^ The diazeniumdiolate was synthesized with the addition of isoamyl nitrite in the presence of ammonia and the compound was purified following an acid-base workup. (**Figure 2B a**) However, this procedure was unsuccessful for glydiazen (**4**) and aladiazen (**5**) (also known as nitrosofungin)^39^ due to the instability of the intermediates. For these compounds, another strategy was developed using a one pot procedure. 2-Nitroethanol was reduced with Pd/C, the reaction was filtered and isoamyl nitrite was directly added. Glydiazen (**4**) and aladiazen (**5**) were obtained after purification using an anion exchange SPE column (TMA-acetate, **Figure 2B b**).

The five newly synthesized derivatives were tested for their ability to induce the *mbo* promoter. Leudiazen (**2**) possessed the highest activity and we therefore hypothesized that it could be the signaling molecule produced by the *mgo* gene cluster (**Figure 2C**). Furthermore, we investigated the activity of the (*R*) and (*S*) enantiomer of leudiazen and no significant difference in potency were observed (**Figure S1**). We decided to continue our study with leudiazen ((*S*)-**2**), which can be synthesized from commercially available (*S*)-leucinol.

### Detection of leudiazen in a Pss culture supernatant extract

In order to increase the sensitivity of our strategy, we developed a UHPLC-MS/MS detection method targeting the predicted signaling molecule leudiazen (**2**) by selective reaction monitoring (SRM). We extracted 2 L of Pss culture supernatant following a modified version of our previously described protocol.^27^ The extract was immediately analyzed to avoid the decomposition of the signaling compound. Using this sensitive UHPLC-MS/MS strategy, we were able to detect and quantified the concentration of leudiazen (**2**) in the SRM mode using the characteristic MS/MS fragmentation pattern of the compound (163 Da to 83 Da). Leudiazen (**2**) was detected in the crude extract and its concentration was determined to be 74 ± 7 nM in the Pss supernatant. No leudiazen (**2**) was detected in the extract of the *ΔmgoA* mutant (**Figure 2D**).

We determined the response of the promoter to different concentrations of leudiazen (**2**). We observed that the addition of synthetic leudiazen ((*S*)-**2**) to the growth medium activated the biosensor strain in a dose dependent manner (**Figure 3A**). Addition of 1 μM and above of leudiazen (**2**) restored the β-galactosidase activity even above the level observed with the sensor plasmid in the wild type background.

**Figure 3.**
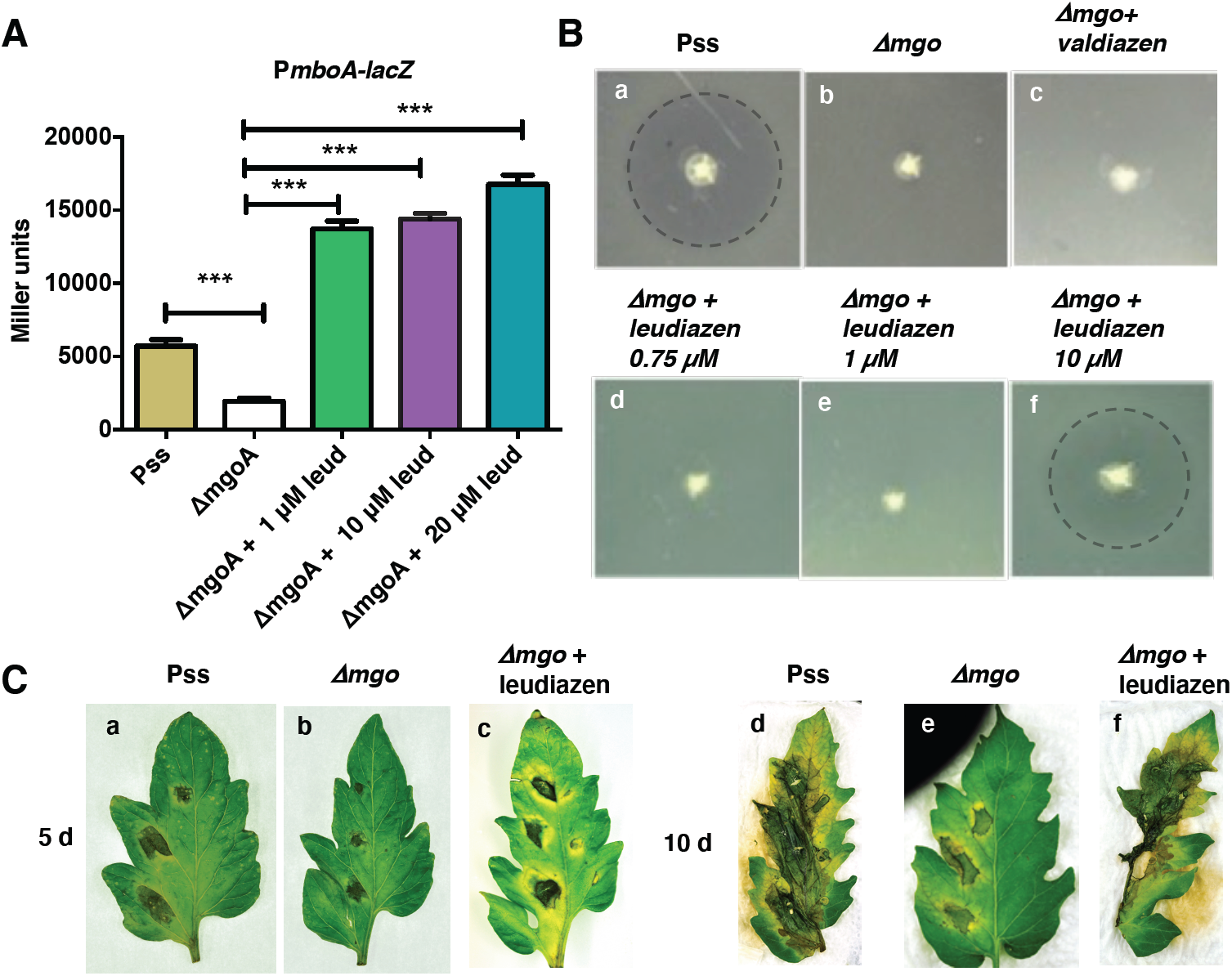
**A)** Activities of the *mboA* promoter in the presence of various concentrations of leudiazen (**2**, leud): The biosensor was grown in media supplemented with various concentrations of synthetic leudiazen ((*S)*-**2**) and β-galactosidase activities were recorded. Results are presented as mean and error bars represent SE. Statistical analysis was performed using one-way ANOVA and Tukey’s multiple comparison as post test. ns, not significant. Significance is indicated by one or three stars. \*\*\**p* < 0.001, *n* = 3; **B)**. Mangotoxin production assay using *E. coli* as an indicator (**a**) Halo around Pss indicates mangotoxin production while there’s no halo around (**b**) *Δmgo* (**c**) *Δmgo* supplemented with valdiazen (**1**); (**d**) *Δmgo* supplemented with 0.75 μM and (**e**) 1 μM leudiazen ((*S)*-**2**). Mangotoxin production was restored when (**f**) *Δmgo* supplemented with 10 μM leudiazen ((*S)*-**2**). The halos are highlighted with a dashed line; **C**). Leudiazen controls virulence of Pss. Tomato leaves were infected with the wild type and the *ΔmgoA* mutant in the absence or presence of 10 μM leudiazen ((*S)*-**2**). Pictures were taken 5 and 10 days post-infection (dpi).

We next tested whether leudiazen (**2**) could restore mangotoxin production of the *ΔmgoA* mutant using an *E. coli* indicator strain. We grew the wild type and the *ΔmgoA* mutant in the absence or presence of different amounts of leudiazen ((*S*)-**2**) and tested for growth inhibition of the *E. coli* indicator strain. In this assay, mangotoxin production is indicated by a halo around the colony. While the *ΔmgoA* mutant did not exhibit a clearing zone (**Figure 3B a**), mangotoxin production was restored when the mutant was grown in the presence of 10 μM leudiazen ((*S*)-**2**) (**Figure 3B f**). The addition of valdiazen (**1, Figure 3B c**) did not rescue the mutant phenotype.

### Leudiazen controls Virulence of Pss

Given that mangotoxin is a major virulence factor of Pss, we examined whether leudiazen (**2**) would also restore virulence of the *ΔmgoA* mutant strain using detached tomato leaflets as infection model. While the wild type strain caused severe necrosis 10 days post infection (dpi) the *ΔmgoA* was strongly attenuated (**Figure 3C**). However, virulence was fully restored when the *ΔmgoA* mutant was grown in the presence of 10 μM leudiazen ((*S)*-**2**).

### Leudiazen is a Volatile Secondary Metabolite

Intrigued by the difficulties we encountered during the isolation of leudiazen (**2**), we decided to investigate the stability of the compound. The degradation of leudiazen ((*S*)-**2**) was studied by ^1^H-NMR in two different solvents, D_2_O and MeOD, with seven times points, 10 min, 60 min, 3 h, 24 h, 48 h, 72 h, 144 h, and at room temperature. Leudiazen (**2**) was found to be stable in MeOD for 48h and in D_2_O for 3 h. Thereafter, the appearance of a by-product was observed (**Figure S3** and **S4**). The partial degradation of leudiazen (**2**) could not explain the loss of the compound in our previous isolation attempts and for this reason we investigated the volatility of the compound. Leudiazen ((*S*)-**2**) was evaporated under a gentle flow of nitrogen or under vacuum, which are the standard procedures we used during our isolation strategy. The weight of the sample was determined over a time period of 8 h. After 4h, we observed that more than 50% of the sample weight was lost and after 8 h most of the compound had evaporated (**Figure S5**). Furthermore, a head-space experiment was performed by heating leudiazen ((*S*)-**2**) in a vial to 100°C for 10 min, the gas phase was transferred into another vial containing MeCN and this solution was analyzed by HRMS (**Figure S6**). The results showed that leudiazen (**2**) was present in the gas phase of the first vial, as it was detected in the MeCN solution. Additionally, the ability of leudiazen (**2**) to diffuse through air was tested on a split plate system using *E. coli* as an indicator strain to assess mangotoxin production. In this experiment, a halo around the colony was an indication of the mangotoxin production by the *mbo* gene cluster which was activated by leudiazen (**2**). While the *ΔmgoA* mutant did not exhibit any antibacterial activity, the presence of leudiazen (**2**) in the adjacent compartment enabled the mutant to restore mangotoxin production and a clearing halo could be observed (**Figure 4 A**). Furthermore, no halo was observed when charcoal, a volatile trap, was added to one of the compartments (**Figure S8**).

**Figure 4.**
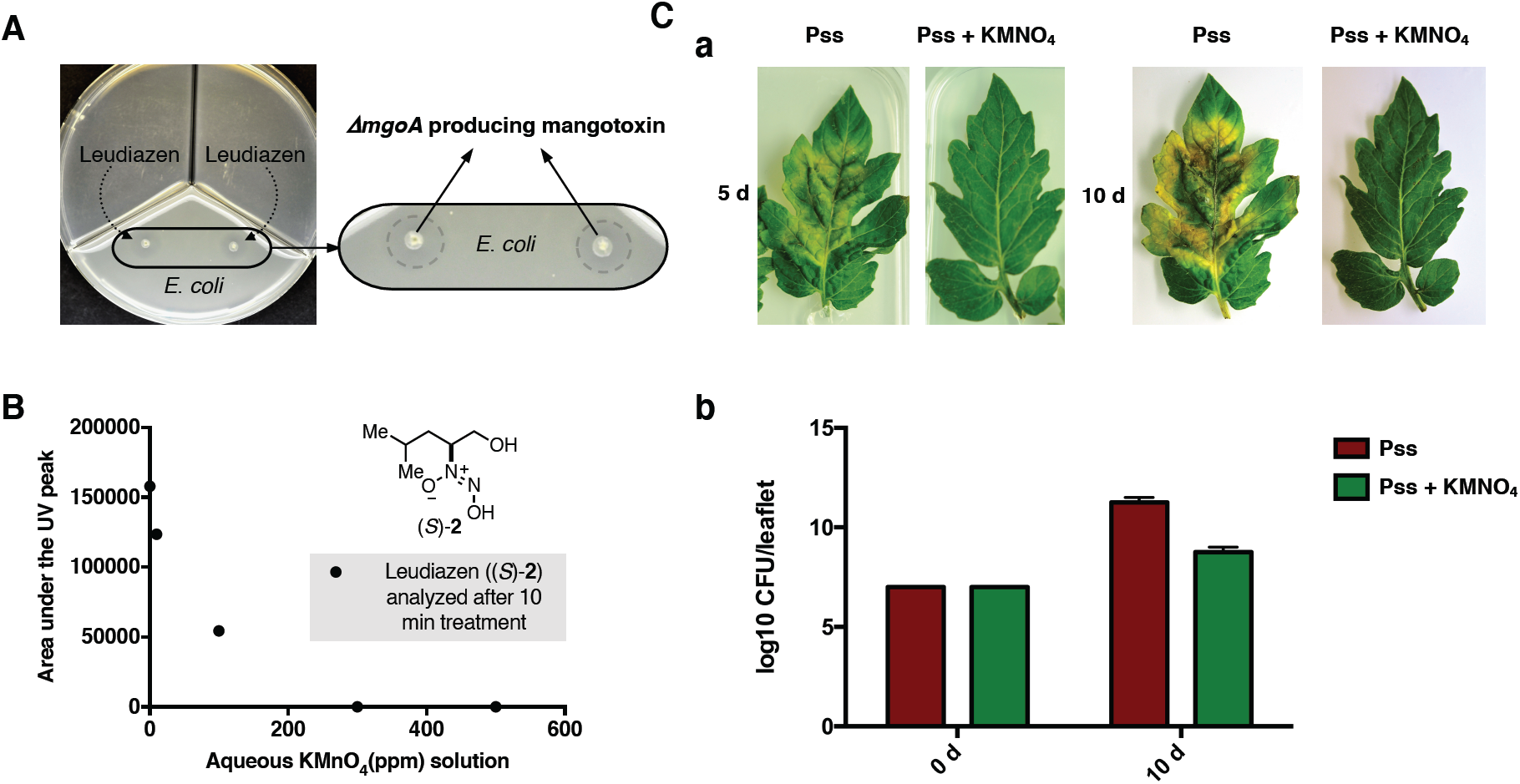
**A)** Split plate assay with leudiazen (**2**) in the adjacent compartments and Δ*mgo* stabbed on two spots over the lawn of *E. coli*. Halos around the Δ*mgo* denotes growth inhibition of *E. coli* by mangotoxin production **B)**. A solution of leudiazen ((*S)*-**2**, 10 μg/mL) was treated with KMnO_4_ at concentrations varying from 09 to 500 ppm. The solutions were stirred for 10 min and analyzed by UHPLC-UV (254 nm). The areas under the UV peaks are reported; **C)** An aqueous KMnO_4_ solution (1000 ppm) protects tomato leaves from *P. syringae* infection. (**a**) Pictures showing tomato leaves infected with the wild type strain (first image: 5 dpi, third image: 10 dpi) and the wild type strain followed by treatment with KMnO_4_ (second image: 5 dpi, fourth image: 10 dpi). (**b**) Amount of *P. syringae* cells recovered at 10 dpi. The red bars represent the recovered bacteria from the untreated leaves and the green plots represent the recovered bacteria from the leaves treated with KMnO_4_.

### Inactivation of Leudiazen

Given that leudiazen controls the virulence of Pss, we hypothesized that inhibition of this communication system could be an effective way to control the pathogenicity of the strain. We therefore sought of developing a chemical strategy for the degradation of leudiazen (**2**) and potentially other diazeniumdiolates. Given that previous work demonstrated the ability of KMnO_4_ to efficiently decompose the diazeniumdiolate functional group,^40^ we treated leudiazen ((*S)*-**2**) with KMnO_4_ at different concentrations (0, 10, 100, 300 and 500 ppm), stirred the solution for 10 min and determined the degradation of the signaling molecule by UHPLC-UV. These experiments revealed that a concentration of 300 ppm of KMnO_4_ was sufficient to completely inactivate leudiazen ((*S)*-**2**) after 10 min **(Figure 4B**).

### Dilute KMnO_4_ solution protects detached tomato leaves from *P. syringae* infection

Since KMnO_4_ treatment efficiently inactivates the leudiazen signal compound, we next tested whether dilute KMnO_4_ solution could protect tomato leaves from infection with Pss. Our results demonstrate that the infected plant leaves that were treated with an aqueous KMnO_4_ solution (1000 ppm) did not possess necrotic symptoms while untreated leaves were severely affected (**Figure 4C a)**. We could recover Pss cells from treated and the untreated leaves (**Figure 4C b**), indicating that the treatment is not affecting bacterial growth.

## Discussion

Pss, a major pathovar among the 60 members of the heterogenous group of *P. syringae*, causes bacterial apical necrosis (BAN) of mango trees resulting in severe economic losses.^41^ Previous studies, based on phenotypic and genetic analysis, revealed that Pss strains isolated from mango constitute a cluster characterized by mangotoxin production and are distinct from the Pss isolates of other hosts.^42^ Moreover, the gene cluster directing the biosynthesis of mangotoxin has been identified ^16^ and mangotoxin production was shown to be controlled by a self-produced signaling molecule.^20,21^ However, neither the structure of the toxin nor of the signal has been elucidated. In our study, we identified this key regulatory signal molecule of Pss named leudiazen (**2**), which is the second representative of the recently described diazeniumdiolate signal family.^27^

Our first isolation attempt was based on a classical bioassay-guided fractionation approach. Pss extract was separated by several analytical techniques and the fractions were analyzed using our reporter strain. We were surprised to observe a loss in biological activity during the fractionation process and we hypothesized that the compound decomposed. Based on the similarities of the *mgoA* NRPS adenylation domain with its homologs,^37^ the signaling molecule produced by Pss was proposed to be an analog of valdiazen (**1**). We embarked on the development of a structure-activity relationship (SAR) of **1**. The structure of the derivatives was designed by replacing the valine moiety of **1** by alanine, leucine, isoleucine, and glycine. Two synthetic strategies were elaborated for the formation of the hydroxylamine key intermediate. Using these approaches, we obtained aladiazen (**5**), leudiazen (**2**), isoleudiazen (**3**), and glydiazen (**4**). The compounds were tested for their ability to rescue *mbo* promoter activity and mangotoxin production in the *ΔmgoA* mutant strain and leudiazen (**2**) was the most effective compound. This result indicated that it may be the indigenous signal produced by Pss (Figure **2C**) and is also in line with a bioinformatics analysis of the substrate specificity of the adenylation domain of MgoA.^37^

To increase the sensitivity of our detection strategy, we developed a UHPLC-MS/MS method using our synthetic standard. An extraction was performed on two liters culture media containing Pss and its *ΔmgoA* mutant. Leudiazen (**2**) was unambiguously identified only in the extract of Pss and the concentration of leudiazen (**2**) was determined to be 74 ± 7 nM in the supernatant. Unfortunately, the absolute configuration of the naturally produced leudiazen (**2**) could not be identified due to the low amount produced by the strain. We hypothesized that leudiazen (**2**) is produced as a racemate as it was shown for valdiazen (**1**). Both enantiomers of leudiazen, ((*S*)-**2**) and ((*R*)-**2**) were evaluated and no significant difference in potency was observed (Figure **S1**). Therefore, the naturally derived enantiomer (*S)*-**2** was used for all the experiments. Interestingly, leudiazen (**2**) concentration was found to be approximately 300 times lower compared to the concentration of valdiazen (**1**) that we recovered from the supernatant of *B. cenocepacia* H111. It is also important to note that we could only restore mangotoxin production of the *ΔmgoA* mutant when the strain was exposed to at least 10 μM leudiazen ((*S*)-**2**). This may indicate that leudiazen (**2**) is either unstable or is not efficiently taken up by the cell or that the organism produces much higher amounts of the signaling molecule under *in vivo* conditions compared to the laboratory growth conditions used. At present we favor the last option, as we observed that the *mboA* promoter activity was induced in the presence of a mango leaf extract (data not shown), suggesting that a plant compound may stimulate leudiazen (**2**) biosynthesis. However, additional work will be required to unravel the spatial and temporal production of leudiazen (**2**) (and consequently mangotoxin) during the infection process.

The chemical properties of leudiazen (**2**) were studied and the results of our experiments suggested that leudiazen (**2**) is volatile (**Figure 4A**). The volatility is a striking feature that differentiates leudiazen (**2**) from other signaling molecules. Very few volatile signals in bacteria have been described to date and they include indole, which is produced by *E. coli* and many other bacteria, as well as methyl 3-hydroxypalmitate (3-OH PAME) and methyl 3-hydroxymyristate ((*R*)-3-OH MAME) by *Ralstonia solanacearum* (*R. solanacearum*).^43–45^ Notably, 3-OH PAME and (*R*)-3-OH MAME, like leudiazen (**2**), control virulence in a major plant pathogen *R. solanacearum*.^43,45^ Volatile signals could be key to long distance communication and may thus be instrumental in establishing and disseminating the infection in the host. Work currently under progress therefore aims at detecting leudiazen (**2**) in the headspace of Pss cultures as well as in the gas phase of plates containing infected tomato leaves to confirm the occurrence of leudiazen as a volatile messenger even in natural conditions.

Most pathogenic bacteria utilize signaling molecules to coordinate the expression of virulence factors, such that the infected host is unable to launch a timely immune response. Hence, interference with bacterial signaling has emerged as a highly valuable strategy for the development of novel antiinfective drugs that do not aim at killing the pathogen but attenuate their virulence.^46^ We exploited the instability of leudiazen (**2**) to develop such an antivirulence strategy. Given that leudiazen-mediated signaling is a major regulator of virulence, we reasoned that degradation of leudiazen (**2**) would impair the pathogenicity of Pss without affecting the viability of the organism. As a consequence, such a treatment regime may be less vulnerable to the appearance of resistant mutants.

In fact quorum sensing inhibition strategies have been extensively studied in different bacteria and most of them described to date involve enzymatic degradation and inactivation of regulatory signals.^47,48^ The potential of employing such signal inhibitors in disease prevention and treatment of *Erwinia carotovora* infection has been shown in various plant models.^49,50^ However, an antivirulence strategy by chemical inactivation of signaling molecule has not been demonstrated. In accordance with a previous study which reported that KMnO_4_ efficiently oxidizes the diazeniumdiolate cupferron,^51^ we provided evidence that it also decomposes leudiazen (**2**) at a concentration as low as 300 ppm (**Figure 4B**) in 10 min. More importantly, the virulence assays reported above underline the strategy that a KMnO_4_ treatment, using a solution concentrated at 1000 ppm, efficiently protects the plant leaves from Pss infection. In this context it is noteworthy that KMnO_4_ displays low toxicity and is used for the treatment of various skin lesions or diseases, including impetigo, pemphigus, dermatitis, and tropical ulcers^52^ as well as an antifungal agent to treat mildew on plant leaves.^53^ Interestingly, KMnO_4_ solutions are regulatorily approved for organic farming for fruit and olive trees in the European Union.^54^

Our results demonstrate that chemical inactivation of leudiazen prevents Pss to produce mangotoxin, a main virulence factor of this organism. Additional work will be required to evaluate the applicability of this approach under field conditions. Indeed, none of the conventional Bordeaux mixture treatments conferred complete protection against BAN in previous field trials^55^ and an effective treatment strategy is urgently required to limit the economic loss due to Pss infections.

The discovery of leudiazen (**2**) confirmed the existence of a new class of signaling molecule composed of diazeniumdiolates compounds. It is particularly interesting that variants of the diazeniumdiolate class of signal molecules regulate different metabolites in phylogenetically diverse bacteria ranging from plant beneficial strains to plant and human pathogens.^21,23,27,56^ This suggests that this novel family of signal molecules controls various phenotypic traits and our future work aims at characterizing the role of these communication systems in diverse bacteria.

## Conclusion

A novel signaling molecule, named leudiazen (**2**), was isolated from the plant pathogen Pss. This natural product belongs to the rare family of compounds possessing a diazeniumdiolate and represents the second member of a new class of signaling molecules. Leudiazen (**2**) was found to be essential for the virulence of the pathogen by the positive regulation of mangotoxin. Additionally, leudiazen (**2**) was found to be volatile using a head-space experiment and was found to decompose in aq. solution. Taking advantage of the instability of leudiazen (**2**), we developed a biocontrol strategy on the basis of dilute KMnO_4_ solution. Leudiazen (**2**) is shown to be sensitive to the treatment with a KMnO_4_ solution and preliminary results from pathogenicity assays indicated that this solution inactivates the signaling molecule without affecting growth of the bacteria. This finding represents the first example of chemical inactivation of a bacterial signaling molecule to attenuate the virulence of a pathogen. Furthermore, our new approach using a KMnO_4_ solution to limit BAN is regulatorily approved for organic farming in the European Union.

## Supporting information

SupInfo

## Author Contributions

All authors contributed to the redaction of the manuscript. The experiments were planned by S. S., A. M., L. E. and K. G. ^‡^The authors S. S. and A. M. contributed equally.

## Funding Sources

The research was funded in part by the Swiss National Science Foundation (CRSII5_186410 and 200021_182043)

## Notes

The authors declare no competing financial interest.

## ACKNOWLEDGMENT

We thank Dr. Kirsty Agnoli for her help with the signal extraction procedures and Dr. Carlotta Fabbri for the excellent technical assistance

## Notes

### Competing Interest Statement

The authors have declared no competing interest.

