## Supplementary material for "A Volatile Signal Controls Virulence in the Plant Pathogen *Pseudomonas syringae* pv. *syringae* and a Strategy for Infection Control in Organic Farming": SupInfo

### 1 General information, materials and equipment

All chemicals were purchased from Acros, Fluka or Sigma-Aldrich and were used without further purification. All reactions were carried out in heat gun-dried glassware (unless aqueous reagents were used) and reactions involving air sensitive compounds were performed under an argon or nitrogen atmosphere. Solvents applied for chemical transformations were either puriss quality or HPLC grade solvents, which had been dried by filtration through activated aluminum oxide under nitrogen ($H_2O$ content <10 ppm, *Karl-Fischer* titration). For work-up and purification solvents were distilled from technical grade. All synthetic transformations were monitored either by thin layer chromatography (TLC) or $^1$H NMR spectroscopy. Yields refer to purified, dried and spectroscopically pure compounds. TLC was performed on Merck silica gel 60 F254 plates (0.25 mm thickness) pre-coated with a fluorescent indicator. Concentration under reduced pressure was performed by rotary evaporation at 40°C. Flash chromatography was performed using silica gel 60 (230-400 mesh) from Sigma-Aldrich with a forced flow eluent at 0.1-0.3 bar pressure. All $^1$H and $^{13}$C NMR spectra were recorded using a Bruker Avance 400 MHz, 500 MHz, 600 MHz ($^1$H) & 101 MHz or 126 MHz ($^{13}$C) spectrometer at RT (unless otherwise stated). Chemical shifts (δ-values) are reported in ppm, spectra were calibrated relative to the residual proton chemical shifts (MeOD, δ = 3.31; $D_2O$, δ = 4.79) and the residual carbon chemical shifts (MeOD, δ = 49.0) of the solvents, multiplicity is reported as follows: s = singlet, d = doublet, t = triplet, q = quartet, m = multiplet or unresolved and coupling constant *J* in Hz. IR spectra were recorded on a Perkin Elmer SpectrumTwo ATR-FTIR. The absorptions are reported in $cm^{-1}$. All mass spectra (ESI-HRMS) were recorded on a maXis instrument (Bruker Daltonics GmbH, Bremen, Germany) equipped with an Apollo electrospray ionization (ESI) source. The mass analyzer was calibrated between *m/z* 118 and 2721 using an *Agilent* ESI-L low concentration tuning mix solution (*Agilent*, USA) at a resolution of 20'000 and a mass accuracy better than 2 ppm. Optical rotations $°[\alpha]_D^T$ were measured at the sodium D line using a 1 mL cell with a 1 dm or 0.1 dm path length on a Jasco P-2000 digital polarimeter and the concentration c is given in g/100 mL $CHCl_3$.

### 2 Microbiological methods

#### 2.1 Bacterial strains, plasmids and growth conditions

Strains and plasmids used in this study are listed in **Table S1**. *Pseudomonas syringae* pv. *syringae* (Pss) wild type UMAF0158, Pss *ΔmgoA* mutant as well as strains carrying transcriptional *lacZ* fusions, Pss pMP/P*mboA-lacZ* and *ΔmgoA* pMP/P*mboA-lacZ* strains were kindly provided by Víctor J. Carrión and Francisco M. Cazorla. The construction of both Pss *ΔmgoA* mutant and the transcriptional fusions has been reported previously.[1,2] *Escherichia coli* (*E. coli*) strains were routinely grown in LB medium (BD Difco, USA) at 37 °C while *P. syringae* strains were grown at 30° C. For the extraction of leudiazen (**2**), as well as for promoter activity measurements, *P. syringae* strains were grown in PMS minimal medium (consists of 1g of $(NH_4)H_2PO_4$ per liter, 0.2 g of KCl per liter, and 0.2 g of anhydrous $MgSO_4$ per liter.)[3] The pH was adjusted to 7.0 before autoclaving, and 0.2% glucose was added as a carbon source. Yeast extract (0.005%) was routinely added to the PMS medium to boost growth. When required, media were supplemented with antibiotics at the following concentrations: kanamycin (Km) at 100 µg/ml, gentamicin (Gm) at 20 µg/ml, tetracycline (Tc) at 20 µg/ml and chloramphenicol (Cm) at 20 µg/ml.

**Table S1**: List of strains used in this study

| **Strains** | **Genotype/characteristics** | **Source** |
|---|---|---|
| ***E. coli*** | | |
| *E. coli* K-12 | F- λ- rph-1 *ilvG- rfb*-50; MG1655 | Eberl lab |
| ***Pseudomonas syringae* pv. *syringae*** | | |
| UMAF0158 | Wild type, isolated from mango, mangotoxin producer, $Nf^r$ | 4 |
| ΔmgoA | *mgoA* deletion mutant of UMAF0158 | 1 |
| *P. syringae* pMP/PmboA-lacZ | Transcriptional *lacZ* fusion of *mbo* promoter in Pss wild type | 2 |
| ΔmgoA pMP/PmboA-lacZ | Transcriptional *lacZ* fusion of *mbo* promoter in Δ*mgoA* mutant | 2 |

### 2.2 Assessment of promoter activity by β-galactosidase assay

Promoter activity of the transcriptional *lacZ* fusions in liquid cultures was assessed by β-galactosidase assays as described before with minor modifications.[5] Briefly, bacterial cells were grown overnight in PMS minimal medium. Where indicated, synthetic valdiazen (**1**) or leudiazen (**2**) or ((S)-**2**) or ((R)-**2**) and different valdiazen derivatives dissolved in methanol were added to the cultures. 50–200 µl of cells were harvested, resuspended in Z-buffer (1 mL) and $OD_{600}$ values were recorded. To permeabilize the cell membrane chloroform (25 µL) and (0.1%) were added to the residual bacterial suspension (1 mL), vortexed for 10 s and incubated at 28 °C for 10 min. The reaction was initiated by adding *o*-nitrophenyl-β-D-galactoside (ONPG, 200 µL) solution (4 mg/ml in Z buffer) to each sample, vortexed briefly and incubated at RT. The reaction was stopped by the addition of an aq. $Na_2CO_3$ soln. (500 µL, 1M). The samples were centrifuged at 16,000 rpm for 10 min and cell debris-free supernatant (1 mL) was used to measure the absorbance at 420 nm and 550 nm. The promoter activity (expressed as Miller units) was determined using the following formula:

$$1 \text{ Miller Unit} = 1000 * \frac{OD_{420} - 1.75 * OD_{550}}{t * v * OD_{600}}$$

Where : t = reaction time in minutes and v = volume of assayed sample in milliliters

Data were based on three independent biological replicates (n = 3). For every β-galactosidase assay, a sample containing only the growth medium was processed as described above and used as a blank.

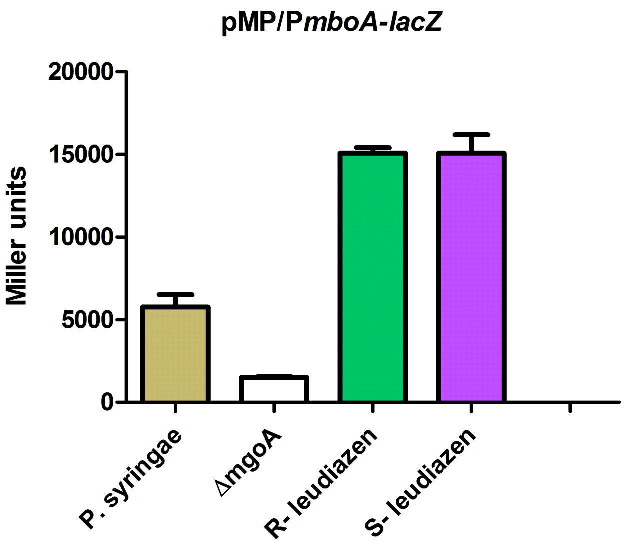


**Figure S1**: The activity of the *PmboA-lacZ* fusion in the presence of (*R*)-**2** and (*S*)-**2** enantiomers of leudiazen. The β-galactosidase activity of the biosensor plasmid was lower in the *ΔmgoA* mutant compared to the activity in the wild type. Both leudiazen enantiomers were found to restore the activity of the *ΔmgoA*. Results are presented as mean and error bars represent SE.

### 2.3 Mangotoxin production assay

Mangotoxin production was determined using an *E. coli* strain as indicator using a previously described procedure[6] with minor modifications. Briefly, a double layer of the indicator microorganism, an *E. coli* K12 strain was prepared and after solidification, the Pss strains to be tested were stab-inoculated (Pss wild type, *Δmgo* mutant and *Δmgo* mutant complemented with different amounts of leudiazen (**2**) or ((S)-**2**) or ((R)-**2**)) on to the agar seeded with *E. coli*. The plates were incubated at 30°C for 48 h and observed for inhibition zones around the colony. *E. coli* killing assay was also performed on split petri dishes to confirm the volatile-mediated effect of leudiazen. PMS medium mixed with *E. coli* ($10^7$ CFU final concentration) was included in one compartment of the petri dish and the *ΔmgoA* mutant was stab inoculated on to it while 10 µl of 20 µM leudiazen was spotted in the adjacent compartments. A control plate, containing charcoal in a 3rd compartment to trap the volatile leudiazen, was also included in the assay (**Figure S2**).

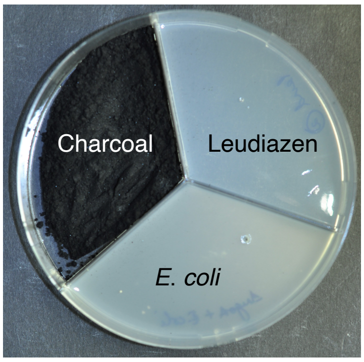


**Figure S2** Split plate assay including charcoal in one compartment which traps the volatile leudiazen (**2**) and avoids mangotoxin production.

### 2.4 Virulence assay using tomato leaflets as model host

To assess whether the putative signal molecules restored the virulence in the mutant strain, detached tomato leaflets (Hellfrucht Frühstamm variety) were used as model hosts and the virulence assay was performed as previously described[6] with some modifications. Exponentially growing cultures of Pss in PMS were adjusted to $10^8$ cfu ml$^{-1}$. In one set of virulence assay, 10 µl drops of the bacterial suspension were injected on three different points on one side of a leaflet while three 10 µl drops of an aq. $MgSO_4$ soln. (10 mM) were injected on the other side of the same leaflet as control. Alternatively, in another set of virulence assay, six 10 µl drops of the bacterial suspension were injected on six different points on the same leaflet and detached leaflets inoculated with an aq. $MgSO_4$ soln. (10 mM) were included in all experiments as a control. Six tomato leaflets were used per strain and the infected leaflets were maintained at 22°C and a 16:8-h light: dark photoperiod for a period of 10 days. These experiments were repeated three times. The development and severity of necrosis on the leaflets was determined every 2 days for a period of 10 days. Bacterial strains were retrieved from the infected leaves and colony counts were calculated on several days post inoculation.

### 2.5 $KMnO_4$ treatment of tomato leaves infected with Pss

To test whether $KMnO_4$ protects the tomato leaves from Pss infection, detached tomato leaves infected with Pss, as described above, were treated with an aq. $KMnO_4$ soln. Leaflets (6) infected with Pss were dipped in an aq. $KMnO_4$ soln. (1000 ppm) with manual shaking at RT, allowing a total contact time of around 20 seconds. The solution was decanted, and the leaves were then dipped in fresh sterile distilled water to remove any traces of $KMnO_4$. The tomato leaflets (6) infected with Pss without $KMnO_4$ treatment were included in the assay. Both set of leaflets was maintained for a period of 10 days and development of necrotic symptoms was recorded. Bacterial strains were recovered from both set of leaflets after 10 days and presence of Pss were confirmed by PCR using Pss specific primers.

### 2.6 Sequence alignment

The sequence alignment of MgoA (accession: AGA16733) from Pss, HamD (accession: CDN62030) from *Burkholderia cenocepacia* H1111 and PvfC (accession: Q1IGU3) from *Pseudomonas entomophila* L48. was performed on Genieous (10.2.5) using the MUSCLE algorithm. The pairwise sequence identity was 45.5%.

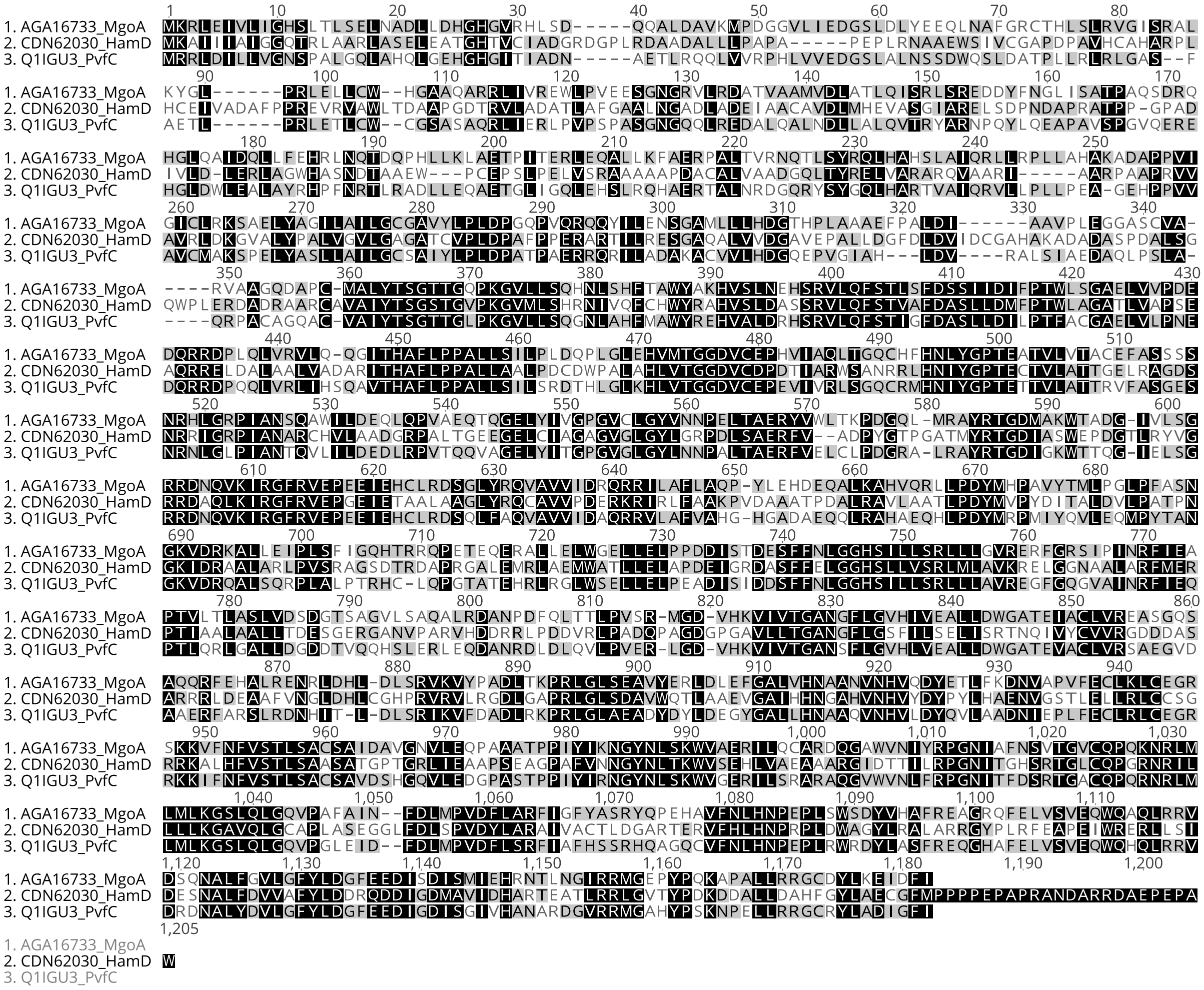


**Figure S3**: Multiple sequence alignment of MgoA (accession: AGA16733) from *Pseudomonas syringae pv. syringae*, HamD (accession: CDN62030) from *Burkholderia cenocepacia* H1111 and PvfC (accession: Q1IGU3) from *Pseudomonas entomophila* L48.

### 3 Analytical methods

#### 3.1 Extraction Procedure

Bacterial strains were grown in PMS minimal medium containing 0.005% yeast extract for 7 days with agitation (220 rpm) at 30 °C. Bacterial cultures were centrifuged for 10 minutes at 8000 rpm in a Sorvall RC-5C plus centrifuge at 4 C. Culture supernatants were subsequently filtered with a Millipore Express$^{Tm}$ Plus 0.22 µm system to remove the cell debris. The cell free supernatant was acidified to a pH of 5 with an aq. HCl soln. (10 M) and extracted thrice with 0.5 volumes dichloromethane. The org. phases were combined, dried with $MgSO_4$, filtered, concentrated under reduced pressure to obtain the crude extract.

#### 3.2 Preliminary purification procedure

##### 3.2.1 Solid phase extraction (SPE)

The crude extract was dissolved in $H_2O$ (1 mL) and passed through SPE (DSC-18, Discovery®, 500 mg), which has been conditioned with MeOH and $H_2O$. A first fraction was obtained by elution with $H_2O$ (6 mL) and a second fraction was obtained by elution with MeOH (6 mL). Each fraction was dried at 35°C over a gentle $N_2$ flow.

##### 3.2.2 Cation and anion exchange columns

The crude extract was separated with the SPE column, as described above, and the dried MeOH fraction was reconstituted in $H_2O$ (1 mL).

Half of the solution (0.5 mL) was treated with a cation exchange SPE column (DSC-SCX, Discovery®, 200 mg), which has been previously conditioned with MeOH and $H_2O$. The $H_2O$ fraction was collected after elution with $H_2O$ (3 mL) and the MeOH fraction after elution with MeOH (2 mL, 2% $NH_4OH$). The $H_2O$ and MeOH fractions were evaporated to dryness.

The second half of the solution (0.5 mL) was treated with an anion exchange SPE column (DSC-SAX, Discovery®, 1 g), which has been previously conditioned with MeOH and $H_2O$. The $H_2O$ fraction was obtained by eluting with $H_2O$ (2mL) and the MeOH fraction by eluting with acidified MeOH (2 mL, 2% $AcO_2H$). The $H_2O$ and MeOH fractions were finally evaporated to dryness.

##### 3.2.3 HPLC separation

The crude extract was fractionated with a SPE column (DSC-18, Discovery®, 500 mg), as described above, the MeOH fraction was separated by HPLC (1100 Agilent system composed of a pump, autosampler, column oven, and UV detector modules) equipped with a RP-HPLC column, Gemini-NX (250 x 4.6 mm, 5 µm) and an variable wavelength detector (254 nm). The solvent system was composed of A, $H_2O$ (0.1% $HCO_2H$) and B, MeCN (0.1% $HCO_2H$), and the flow was 1 ml/min. The gradient was isocratic for 1 min at 2% of B, varied from 2% to 100% of B in 14 min, was kept at 100% of B for 5 min, and finally the column was re-equilibrated at 2% of B for 5 min. Fractions were collected from 0-8, 8-9.6, 9.6-10.5 and 10.5 to 25 min and concentrated to dryness with a gentle flow of $N_2$ gas. The most active fraction collected from 8 to 9 min was further separated using a different solvent system composed of A, $H_2O$ (0.1% $NH_4OAc$), and B, MeCN:$H_2O$ (95:5 with 0.1% $NH_4OAc$). The gradient was identical as described above, the fractions were collected from 0-8, 8-9, 9-10 and 10-25 min, and concentrated to dryness with a gentle flow of $N_2$ gas.

### 3.3 Method for the detection and quantification of leudiazen (2)

Leudiazen (**2**) was quantified by UHPLC-ESI-MS/MS (Ultimate 3000 LC, Thermo Fisher Scientific, coupled to a TSQ Quantum Ultra, Thermo Fisher Scientific) using a Kinetex® EVO C18 (50 x 2.1 mm, 1.7 µm) column at a flow rate of 0.4 ml/min. The solvent system was composed of A ($H_2O$ with 0.1% $HCO_2H$) and B (MeCN with 0.1% $HCO_2H$). After isocratic elution at 5% B for 1 min, the gradient varied from 5% B to 60 % in 2.5 min, 60% to 95% of B in 1 min, 95% to 100% of B in 0.05 min, and the column was finally washed with 100% B for 1.24 min. The detection was achieved in the SRM mode using the specific fragmentation of the protonated molecules $[M+H]^+$ at *m/*z 163 into the fragment ion at *m/z* 83 Da at a collision energy of 14 eV. The quantification was done using a calibration curve obtained from analytical standard solutions prepared in MeOH at the following concentrations: 1, 5, 10, 20, 50, and 100 µg/mL. A sample volume of 3 µL was injected.

#### 3.3.1 Quantification of leudiazen (2) in Pss

Cultures of Pss (2 L) and the *Δmgo* mutant (2 L) were prepared following the extraction procedure described above (**3.1**). The crude extract was resuspended in MeOH (1 mL). A concentration of 23 ± 3 µg/mL of leudiazen (**2**) was determined in the Pss extract using a calibration curve (**Figure S4**), which represents a leudiazen (**2**) concentration of 74 ± 7 nM in the supernatant of Pss. Leudiazen (**2**) was not detected in the *Δmgo* crude extract.

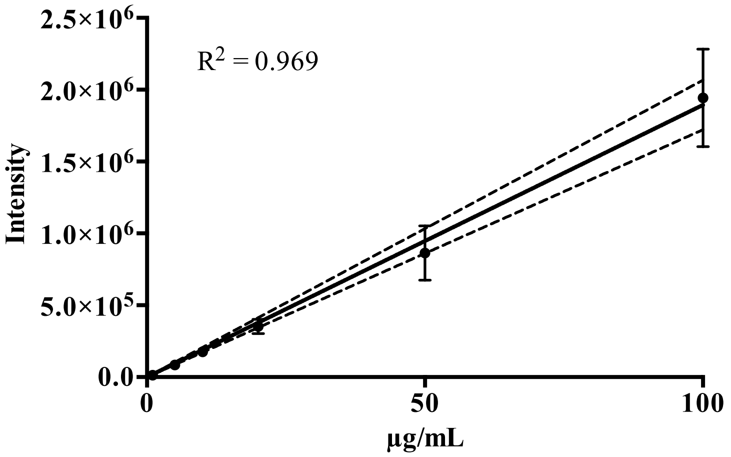


**Figure S4**: Calibration curve for the quantification of leudiazen (**2**)

### 3.4 Degradation procedures

#### 3.4.1 $KMnO_4$ treatment of leudiazen (2)

Aq. $KMnO_4$ soln. (500 µL at a concentration of 20 ppm, 200 ppm, 600 ppm or 1000 ppm) were added to 500 µL of an aq. soln. of leudiazen ((*S*)-**2**, 10 µg/mL) to obtain solutions having a concentration of $KMnO_4$ of 10, 100, 300 and 500 ppm. The samples were stirred at RT for 10 min, filtered and 3 µL injected to the UHPLC-UV system (254 nm).

#### 3.4.2 Degradation study of leudiazen (2) by $^1$H-NMR spectroscopy

Leudiazen ((*S*)-**2,** 10 mg) was dissolved either in $D_2O$ (0.4 – 0.5 mL, **Figure S5**) or in MeOD (0.4 mL – 0.5 mL, **Figure S6**) and both samples were analyzed by $^1$H-NMR after 10 min, 60 min, 3 h, 24 h, 48 h, 72 h, and 144 h.

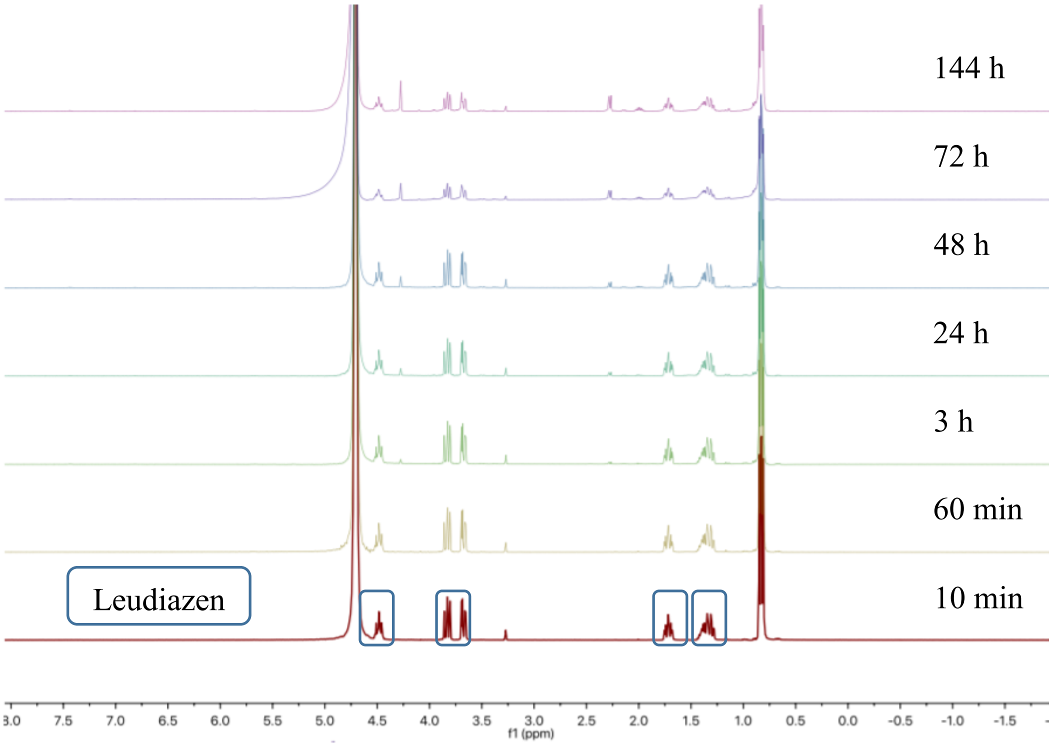


**Figure S5**: Degradation of leudiazen (**2**) in $D_2O$ at RT followed by $^1$H-NMR.

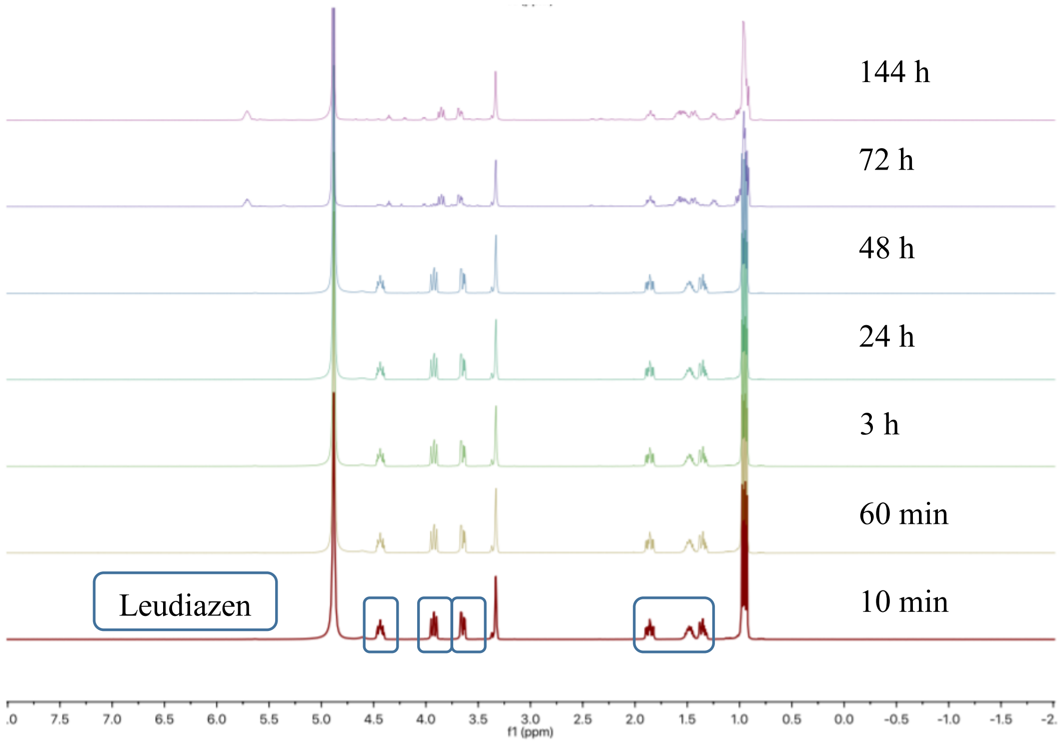


**Figure S6**: Degradation of leudiazen (**2**) in MeOD at RT followed by $^{1}$H-NMR.

### 3.5 Volatility experiment

A glass vial (2 mL) containing leudiazen (((*S*)-**2**), ~5 mg) was placed into a vacuum chamber at 0.02 mbar and another sample was placed under a gentle flow of nitrogen at 40°C. The samples were weighted each 2 h for 8 h (**Figure S7**).

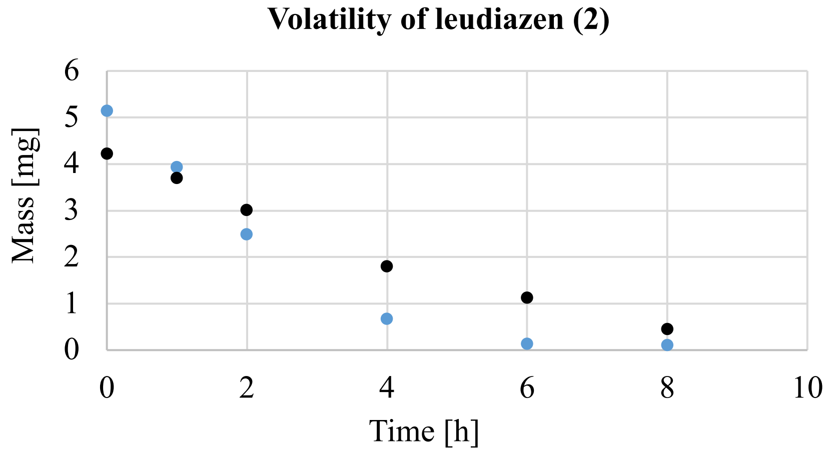


**Figure S7**: Evaporation of leudiazen ((*S*)-**2**) under high vacuum (blue dots) and under gentle flow of nitrogen at 40 °C (black dots).

The headspace experiment was performed by heating leudiazen ((S)-**2**, 0.9 mg) in a 10 mL vial and for 10 min at 100°C (**Figures S8 and S9**). Then, 10 mL of the gas phase was transferred into another vial containing 1 mL acetonitrile with a gas-tight syringe. This vial was shaken for 30 s and an aliquot of the acetonitrile layer (3 µL/min) was measured with ESI-HRMS. ESI-HRMS: *m/z* 161.0930 ($C_6H_{13}N_2O_3^-$ $[M\text{-}H]^-$; calc. 161.0932, -1.2 ppm). A control experiment was performed by injecting a MeCN soln. (**Figure S9**)

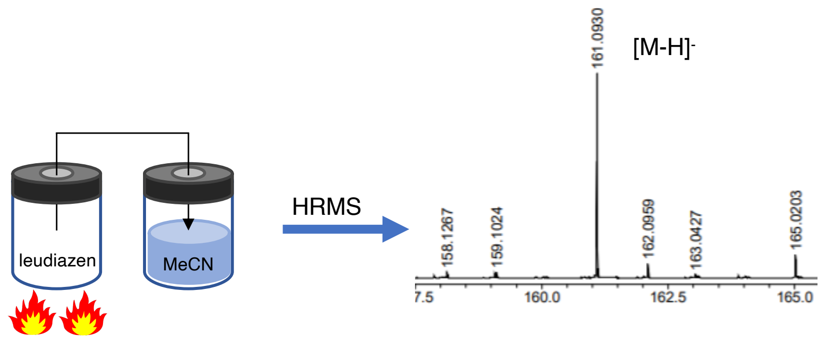


**Figure S8**: Head space experiment to evaluate the volatility of leudiazen (**2**). (*S*)-**2** was heated for 10 min at 100°C and the gas phase was transferred into another vial containing MeCN with a gas tight syringe. Leudiazen (**2**, *m*/*z* 161.093) could be detected by ESI-HRMS in negative mode

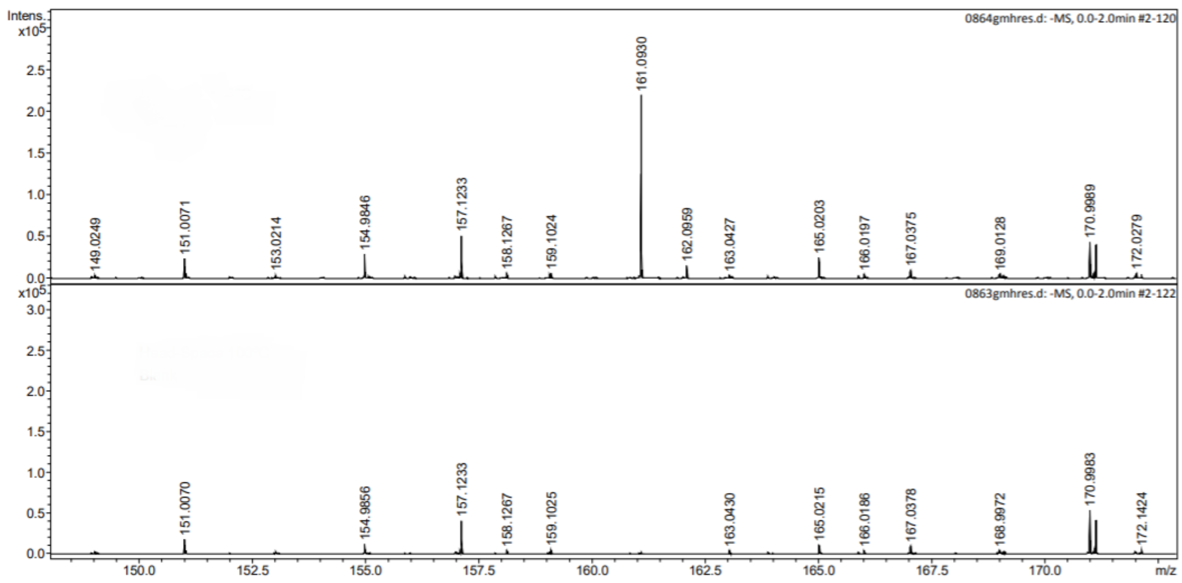


**Figure S9**: Headspace experiment with a 10 mL vial containing leudiazen ((*S*)-**2**, 0.9 mg). Top: 1 mL of the gas phase dissolved in 1 mL MeCN. Bottom: blank control (MeCN).

### 4 Synthesis

#### 4.1 Synthesis of (*S*)-2-(hydroxyamino)-4-methylpentan-1-ol ((*S*)-7)

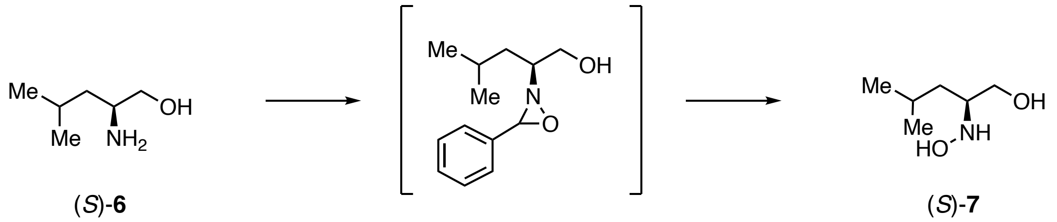


(*S*)-**6** (*S*)-**7**

This experiment was conducted using a modified published procedure.[7] Anhydrous $Na_2CO_3$ (814 mg, 7.7 mmol) and *p*-anisaldehyde (0.64 mL, 5.12 mmol) were added to a soln. of (*S*)-2-amino-4-methylpentan-1-ol ((*S*)-**6**, 600 mg, 5.12 mmol) in dry $CH_2Cl_2$ (20 mL) at RT, and the reaction was stirred at RT for 18 h. The suspension was filtered through a short pad of celite and the solvent was concentrated under reduced pressure to yield a slightly yellowish oil. The oil was dissolved in dry $CH_2Cl_2$ (5 mL) and cooled to 0 °C. A soln. of *m*-CPBA (1.03 g, 4.61 mmol, previously dried over $MgSO_4$ and filtered) in dry $CH_2Cl_2$ (20 mL) was slowly added causing the formation of a white precipitate. The suspension was stirred at 0 °C for 1 h, warmed to RT and stirred for an additional 18 h. The white suspension was filtered off, the residue washed with an aq. sat. $NaHCO_3$ soln. (8 mL) and brine (8 mL), dried over $MgSO_4$ and was concentrated under reduced pressure to yield a yellow oil. The oil was dissolved in dry MeOH (12 mL) and hydroxylamine hydrochloride (712 mg, 7.86 mmol) was added. After 18 h of stirring at RT the reaction mixture was concentrated under reduced pressure. $H_2O$ (12 mL) was added and the aq. layer was then washed with $Et_2O$ (10 x 10 mL), basified with solid $NaHCO_3$ to a pH = 8 and extracted with $Et_2O$ (10 x 10 ml). The combined org. layers were dried over $Na_2SO_4$, filtered, and concentrated under reduced pressure yielding a colourless oil (268 mg). The compound (*S*)-**7** was used in the next step without any further purification.

### 4.2 Synthesis of leudiazen ((*S*)-2)

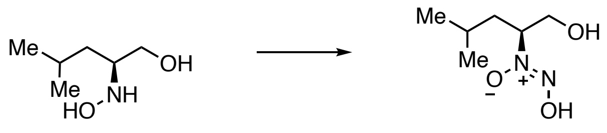


This experiment was conducted using a modified procedure.[7] To a soln. of the hydroxylamine (*S*)-**7** (284 mg) in ammonia soln. (7 N in MeOH, 4 mL) at 0 °C, isoamyl nitrite (0.91 mL, 6.77 mmol) was added dropwise and the reaction mixture was stirred at 0 °C for 30 min. The reaction mixture was warmed to RT and concentrated under reduced pressure. An aq. NaOH soln. (1 M, 1.5 mL) was added and the aq. layer was washed with $Et_2O$ (5 x 3 mL) acidified with an aq. HCl soln. (1 M) to pH = 2 and extracted with $Et_2O$ (5 x 3 mL). The combined org. layers were dried over $MgSO_4$, filtered, and concentrated under reduced pressure. The residue was dissolved in $H_2O$, filtered through a Discovery® DSC-18 SPE (500 mg, supelco) and washed with $H_2O$ (20 mL). The solvent was concentrated under reduced pressure, and the residual $H_2O$ removed by three cycles of dissolving in MeOH (3 x 1.5 mL) followed by concentration under reduced pressure, yielding (*S*)-leudiazen (*S*)-**2** (132 mg, 0.81 mmol, 16% over 4 steps) as a colorless oil.

**Optical rotation**: $[\alpha]_D^{25}$ = –14.6 (*c* 0.84, MeOH); $\mathbf{R_f}$= 0.5 ($CH_2Cl_2$:MeOH, 4:1); **Mp** 76 °C; **FTIR**: ṽ = 3371, 2960, 2874, 1456,1389, 1370, 1345, 1326, 1282, 1237, 1172, 1136, 1062, 992, 936, 672, 559, 501 $cm^{-1}$; $^{1}$**H NMR** (400 MHz, MeOD) $\delta$ = 4.43 (ddt, *J* = 10.6, 9.3, 3.8 Hz, 1H), 3.92 (dd, *J* = 11.8, 9.3 Hz, 1H), 3.65 (dd, *J* = 11.8, 3.9 Hz, 1H), 1.86 (ddd, *J* = 14.0, 10.7, 4.5 Hz, 1H), 1.58–1.41 (m, 1H), 1.35 (ddd, *J* = 14.0, 9.4, 3.8 Hz, 1H), 0.97 (dd, *J* = 6.6 Hz, 3H), 0.94 (d, *J* = 6.6 Hz, 3H); $^{13}$**C-NMR** (101 MHz, MeOD) $\delta$ = 74.9, 63.2, 37.9, 25.9, 23.3, 21.7; **ESI-HRMS** (MeOH): *m/z* 161.0930 ($C_6H_{13}N_2O^-$; $[M-H]^-$; calc. 161.0932).

### 4.3 Synthesis of (*S*)-2-(hydroxyamino)-4-methylpentan-1-ol (7)

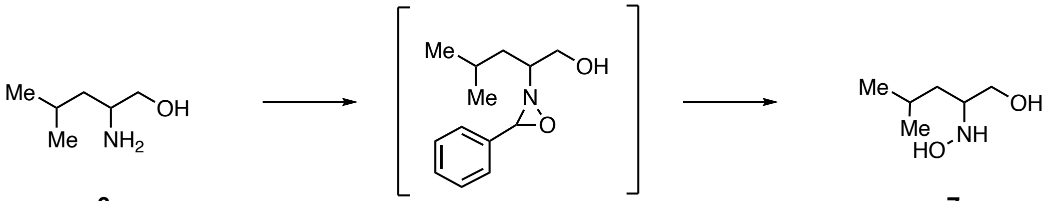


6 7

This experiment was conducted using a modified published procedure.[7] To a soln. of 2-amino-4-methylpentan-1-ol (**6**, 800 mg, 6.83 mmol) in $CH_2Cl_2$ (36 mL), *p*-anisaldehyde (0.85 mL, 6.83 mmol) was added at RT, and the reaction was stirred for 4.5 h, warmed to 40 °C, and stirring was continued for an additional 1.5 h. The soln. was dried over $MgSO_4$, filtered and evaporated under reduced pressure to yield a colourless oil. The oil was dissolved in dry $CH_2Cl_2$ (5.4 mL) and cooled to 0 °C. A soln. of *m*-CPBA (1.68 g, 6.83 mmol, previously dried over $MgSO_4$) in dry $CH_2Cl_2$ (17 mL) was added causing the formation of a white precipitate. The suspension was stirred at 0 °C for 1 h. After 1 h the reaction mixture was warmed to RT and stirred for an additional 1 h. The white suspension was filtered, washed with an aq. sat. $NaHCO_3$ soln. (8 mL) and brine (8 mL), dried over $MgSO_4$ and concentrated under reduced pressure to yield a yellow oil. The oil was dissolved in dry MeOH (8.9 mL) and hydroxylamine hydrochloride (969 mg, 13.7 mmol) was added. After 3 days of stirring at RT the reaction mixture was concentrated under reduced pressure. $H_2O$ (3.4 mL) was added and the aq. layer was then washed with $Et_2O$ (10 x 10 mL), basified with solid $NaHCO_3$ to a pH of 8 and extracted with $Et_2O$ (10 x 10 ml). The combined org. layers were dried over $Na_2SO_4$, filtered, and concentrated under reduced pressure yielding **7** (194 mg) as a colourless oil. The compound **7** was used in the next step without any further purification.

### 4.4 Synthesis of leudiazen (2)

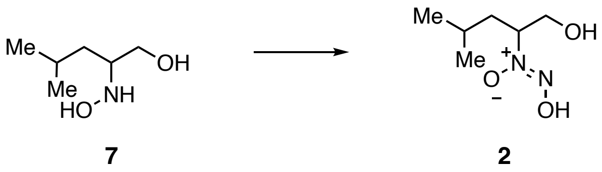

This experiment was conducted using a modified published procedure.[7] To a soln. of the hydroxylamine 7 (420 mg) in MeOH (10 mL) at 0 °C, isoamyl nitrite (1.27 mL, 9.45 mmol) was added dropwise. Ammonia gas was bubbled in the yellowish soln. for 10 min, and the reaction mixture was then warmed to RT and stirred for an additional 20 min. The reaction mixture was concentrated under reduced pressure. Aq. NaOH soln. (1 M, 6 mL) was added and the aq. layer was washed with $Et_2O$ (5 x 15 mL) acidified with an aq. HCl soln. (1 M) to pH = 2 and extracted with $Et_2O$ (5 x 15 mL). The combined org. layers were dried over $Na_2SO_4$, filtered, and concentrated under reduced pressure yielding **2** (349 mg, 2.15 mmol, 15% over 4 steps).

**$^1$H NMR (400 MHz, MeOD)** $\delta$ = 4.42 (ddt, $J$ = 10.6, 9.3, 3.8 Hz, 1H), 3.90 (dd, $J$ = 11.8, 9.3 Hz, 1H), 3.63 (dd, $J$ = 11.8, 3.9 Hz, 1H), 1.84 (ddd, $J$ = 14.1, 10.7, 4.5 Hz, 1H), 1.56–1.39 (m, 1H), 1.33 (ddd, $J$ = 13.9, 9.3, 3.8 Hz, 1H), 0.95 (d, $J$ = 6.5 Hz, 3H), 0.92 (d, $J$ = 6.6 Hz, 3H); **$^{13}$C-NMR (101 MHz, MeOD)** $\delta$ = 74.8, 63.2, 37.9, 25.9, 23.3, 21.7; **ESI-HRMS** (MeOH): $m/z$ 185.0895 ($C_6H_{14}N_2O_3Na^+$; $[M+Na]^+$; calc. 185.0897).

### 4.5 Synthesis of (*R*)-2-(hydroxyamino)-4-methylpentan-1-ol (*R*)-7)

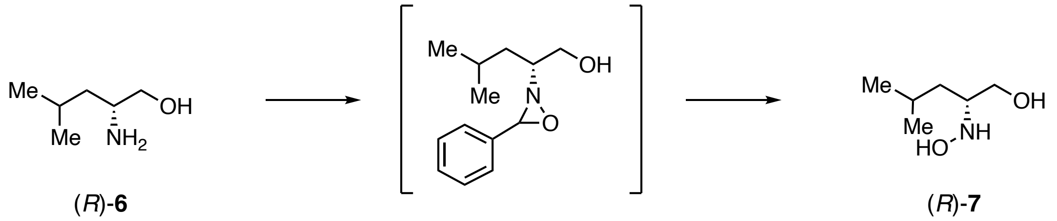


(*R*)-**6** (*R*)-**7**

This experiment was conducted using a modified published procedure.[7] To a soln. of (*R*)-2-amino-4-methylpentan-1-ol ((*R*)-**6**, 1000 mg, 8.53 mmol) in dry $CH_2Cl_2$ (30 mL) anhydrous $Na_2CO_3$ (1360 mg, 12.8 mmol) and *p*-anisaldehyde (1.1 mL, 8.53 mmol) were added at RT, and the reaction was stirred at RT for 2 h. The suspension was filtered through a short pad of celite and the solvents was concentrated under reduced pressure to yield a slightly yellowish oil. The oil was dissolved in dry $CH_2Cl_2$ (10 mL) and cooled to 0 °C. A soln. of *m*-CPBA (1.82 g, 8.10 mmol, previously dried over $MgSO_4$ and filtered) in dry $CH_2Cl_2$ (40 mL) was slowly added causing the formation of a white precipitate. The suspension was stirred at 0 °C for 1 h, warmed to RT and stirred for an additional 1 h. The white suspension was filtered off, the residue was washed with aq. sat. $NaHCO_3$ soln. (8 mL) and brine (8 mL), dried over $MgSO_4$ and concentrated under reduced pressure to yield a yellow oil. The oil was dissolved in dry MeOH (12 mL) and hydroxylamine hydrochloride (1185 mg, 17.10 mmol) was added. After 2 h of stirring at RT. the reaction mixture was concentrated under reduced pressure. $H_2O$ (12 mL) was added and the aq. layer was then washed with $Et_2O$ (10 x 10 mL), basified with solid $NaHCO_3$ to pH = 8 and extracted with $Et_2O$ (10 x 10 ml). The combined org. layers were dried over $Na_2SO_4$, filtered, and concentrated under reduced pressure yielding a colourless oil (223 mg). The compound (*R*)-**7** was used in the next step without any further purification.

### 4.6 Synthesis of leudiazen ((*R*)-2)

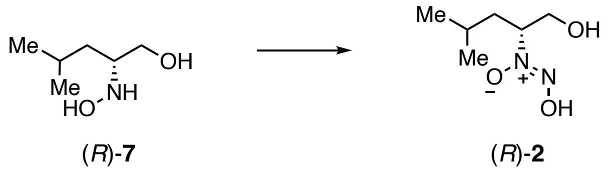

This experiment was conducted using a modified procedure.[7] To a soln. of the hydroxylamine (112 mg) in ammonia soln. (7 N in MeOH, 2 mL) at 0 °C, isoamyl nitrite (0.30 mL, 2.25 mmol) was added dropwise and the reaction mixture was stirred at 0 °C for 30 min. The reaction mixture was warmed to RT and concentrated under reduced pressure. An aq. NaOH soln. (1 M, 1.5 mL) was added and the aq. layer was washed with $Et_2O$ (5 x 3 mL) acidified with an aq. HCl soln. (1 M) to pH = 2 and extracted with $Et_2O$ (5 x 3 mL). The combined org. layers were dried over $MgSO_4$, filtered, and concentrated under reduced pressure. The residue was dissolved in $H_2O$, filtered through a Discovery® DSC-18 SPE (500 mg, supelco) and washed with $H_2O$ (12 mL). The solvent was concentrated under reduced pressure, and the residual $H_2O$ removed by three cycles of dissolving in MeOH (3 x 1.5 mL) followed by concentration under reduced pressure, yielding leudiazen (*R*)-**2** (73 mg, 0.45 mmol, 11 % over 4 steps) as a colorless oil.

**Optical rotation**: $[\alpha]_D^{25}$ = 22.6 (*c* 0.72, MeOH); **$^1$H NMR** (400 MHz, MeOD) $\delta$ = : 4.45–4.38 (*m*, 1 H); 3.90 (*dd*, *J* = 11.7, 9.4 Hz, 1 H); 3.63 (*dd*, *J* = 11.8, 3.8 Hz, 1 H); 1.84 (*ddd*, *J* = 14.0, 10.6, 4.5 Hz, 1 H); 1.52–1.40 (*m*, 1 H); 1.33 (*ddd*, *J* = 14.0, 9.4, 3.8 Hz, 1 H); 0.93 (*dd*, *J* = 12.7, 6.6 Hz, 6 H). **$^{13}$C NMR** (101 MHz, MeOD) $\delta$ = : 74.9; 63.2; 37.9; 25.9; 23.3; 21.7.

### 4.7 Synthesis of glydiazen (4)

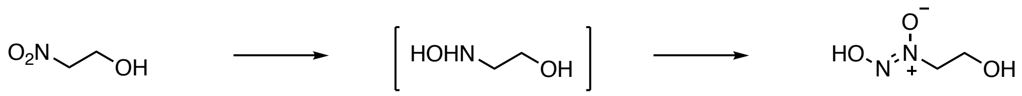

8 4

This experiment was conducted using a modified published procedure.[8] A vigorously stirred soln. of 2-nitroethanol (**8**, 0.29 mL, 4.00 mmol) in degassed EtOH (4 mL) was treated with palladium on charcoal (10% Pd basis, 40 mg) and $H_2$ (1 atm.). The reaction was stirred for 2 h at RT and the reaction mixture was filtered over a pad of celite and washed with MeOH (4 mL). Isoamyl nitrite (1.6 mL, 12.0 mmol) was added dropwise at 0 °C before bubbling ammonia gas through the soln. for 15 min at 0 °C. The mixture was warmed to RT and stirring was continued for 20 min. The reaction mixture was concentrated under reduced pressure and purified by SPE to afford glydiazen (**4**, 77.0 mg, 0.726 mmol, 18% over 2 steps) as a yellowish white solid.

**R*f*** = 0.05 ($CH_2Cl_2$/MeOH, 8:2); **FTIR** : ṽ = 3331, 3029, 2897, 2784, 2475, 2116, 1641, 1551, 1515, 1467, 1428, 1412, 1300, 1283, 1238, 1207, 1098, 1065, 1041, 996, 931, 887, 851, 680, 644, 625, 493, 475 $cm^{-1}$; **$^1$H NMR** (400 MHz, MeOD) $\delta$ = 4.25– 4.13 (m, 2H), 4.06–3.91 (m, 2H);**$^{13}$C NMR** (101 MHz, MeOD) $\delta$ = 65.4, 58.5; **ESI-HRMS** ($H_2O$): *m/z* 129.0270 ($C_2H_6N_2O_3Na^+$; $[M+Na]^+$: calc. 129.0271).

### 4.8 Synthesis of aladiazen (5)

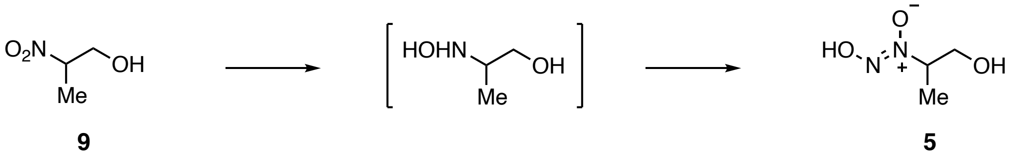


This experiment was conducted using a modified published procedure.[8] To a preheated (45 °C) soln. containing 2-nitropropanol (**9**, 1 mL, 11.3 mmol) and $NH_4Cl$ (1.63 g, 30.5 mmol) in an aq. MeOH soln. (MeOH/$H_2O$ 8:2, 7.3 mL), purified zinc powder (1.61 g, 24.6 mmol, washed with 2% HCl, $H_2O$ and acetone and dried to constant mass) was added in small portions during 10 min so that the temperature rose to 60–65 °C generating a white-grey suspension. The temperature was then kept at 65° for 13 min. Stirring was continued for 15 min during which the suspension cooled to RT. The reaction mixture was filtered and washed with MeOH (15 mL). The soln. was treated with isoamyl nitrite (2.9 mL, 28.4 mmol) at 0 °C and ammonia gas was bubbled for 10 min. The yellow soln. was warmed to RT, stirred for 15 min and concentrated to 1 mL. An aq. NaOH soln. (1 M, 2 mL) was added. The aq. layer was washed with EtOAc (4 x 5 mL) acidified to a pH of 2 with an aq. HCl soln. (1 M) and extracted with $Et_2O$ (12 x 10 mL) and EtOAc (15 x 10 mL) to yield aladiazen (nitrosofungin, **5**, 425 mg, 3.54 mmol, 31% over 2 steps) as a yellowish solid.

**$R_f$** = 0.1 ($CH_2Cl_2$/MeOH, 8:2); **FTIR** $\tilde{\nu}$ = 3188, 3067, 2992, 2976, 2936, 2781, 2521, 1669, 1580, 1455, 1440, 1420, 1385, 1322, 1288, 1265, 1243, 1139, 1116, 1077, 1069, 1030, 947, 897, 860, 680, 637, 515, 504, 493 $cm^{-1}$; **$^1$H NMR** (400 MHz, MeOD) $\delta$ = 4.53–4.36 (m, 1H), 3.90 (dd, $J$ = 11.8, 8.9 Hz, 1H), 3.67 (dd, $J$ = 11.8, 4.0 Hz, 1H), 1.37 (d, $J$ = 6.8 Hz, 3H); **$^{13}$C NMR** (101 MHz, MeOD) $\delta$ = 71.9, 63.6, 14.7; **ESI-HRMS** (MeOH): *m/z* 143.0427 ($C_3H_8N_2O_3Na^+$; $[M+Na]^+$; calc. 143.0427)

#### 4.9 Synthesis of (2*S*,3*S*)-2-(hydroxyamino)-3-methylpentan-1-ol (11)

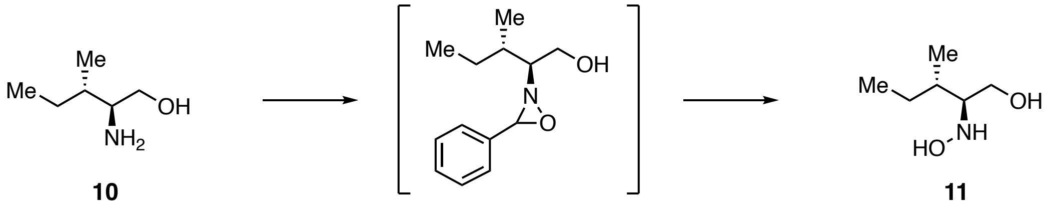


This experiment was conducted using a modified published procedure.[7] To a soln. of (*S*)-(+)-isoleucinol (**10**, 400 mg, 3.41 mmol) in $CH_2Cl_2$ (18 mL), *p*-anisaldehyde (0.42 mL, 3.41 mmol) was added at RT and stirred for 4.5 h and the reaction was warmed to 40 °C and stirred for 1.5 h. The soln. was dried over $MgSO_4$, filtered and evaporated under reduced pressure to yield a colourless oil. The oil was dissolved in dry $CH_2Cl_2$ (2.7 mL) and cooled to 0 °C. A soln. of *m*-CPBA (841 mg, 3.41 mmol, previously dried over $MgSO_4$) in dry $CH_2Cl_2$ (8.5 mL) was added causing the formation of a white precipitate. The suspension was stirred at 0 °C for 1 h. After 1 h the reaction mixture was warmed to RT and stirred for an additional 1 h. The white suspension was filtered, washed with an aq. sat. $NaHCO_3$ soln. (4 mL) and brine (4 mL), dried over $MgSO_4$ and concentrated under reduced pressure to yield a yellow oil. The oil was dissolved in dry MeOH (4.4 mL) and hydroxylamine hydrochloride (484 mg, 6.82 mmol) was added. After 3 days of stirring at RT the reaction mixture was concentrated under reduced pressure, $H_2O$ (3.4 mL) was added and the aq. layer was washed with $Et_2O$ (10 x 5 mL), basified with solid $NaHCO_3$ to a pH = 8 and extracted with $Et_2O$ (10 x 5 ml). The combined org. layers were dried over $Na_2SO_4$, filtered, and concentrated under reduced pressure yielding **11** (194 mg) as a colourless oil. The compound **11** was used in the next without any further purification.

### 4.10 Synthesis of isoleudiazen (3)

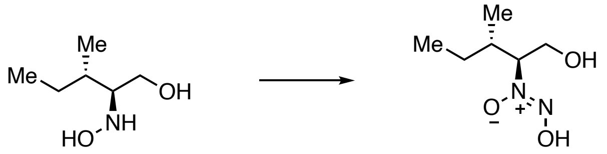


**11** **3**

This experiment was conducted using a modified published procedure.[7] To a soln. of the hydroxylamine **11** (162 mg, 1.22 mmol) in MeOH (5 mL) at 0 °C, isoamyl nitrite (0.51 mL, 3.66 mmol) was added dropwise. To the yellowish soln. ammonia gas was bubbling for 12 min then warmed to RT and the reaction was stirred for an additional 20 min. The reaction mixture was concentrated under reduced pressure. An aq. NaOH soln. (1 M, 3 mL) was added and the aq. layer was washed with $Et_2O$ (5 x 5 mL) acidified with an aq. HCl soln. (1 M) to a pH = 2 and extracted with $Et_2O$ (5 x 5 mL). The combined org. layers were dried over $Na_2SO_4$, filtered, and concentrated under reduced pressure yielding isoleudiazen (**3**, 45.0 mg, 0.277 mmol, 10% over 4 steps) as a yellowish solid.

**$R_f$** = 0.5 ($CH_2Cl_2$:MeOH, 4:1); **Optical rotation**: $[\alpha]_D^{25}$ = −3.18° (*c* 8.8, MeOH); **FTIR**: ṽ = 3369, 3058, 2968, 2938, 2881, 1631, 1456, 1386, 1362, 1319, 1277, 1249, 1212, 1139, 1119, 1058, 1010, 983, 955, 917, 878, 841, 817, 779, 698, 612, 550, 505, 485 $cm^{-1}$; **$^1$H NMR** (400 MHz, MeOD) $\delta$ = 4.12 (td, $J$ = 9.3, 3.3 Hz, 1H), 3.99 (dd, $J$ = 11.7, 9.6 Hz, 1H), 3.83 (dd, $J$ = 11.8, 3.3 Hz, 1H), 2.06–1.83 (m, 1H), 1.48–1.26 (m, 1H), 1.23–1.05 (m, 1H), 0.99 (d, $J$ = 6.9 Hz, 3H), 0.90 (t, $J$ = 7.5 Hz, 3H); **$^{13}$C-NMR** (101 MHz, MeOD) $\delta$ = 81.0, 61.1, 35.3, 26.5, 15.4, 10.8; **ESI-HRMS** (MeOH): *m/z* 185.0896 ($C_6H_{14}N_2O_3Na$; $[M+Na]^+$; calc. 185.0897).

### NMR Spectra

$^1$H-NMR Spectrum of leudiazen ((*S*)-**2**) in MeOD (400 MHz)

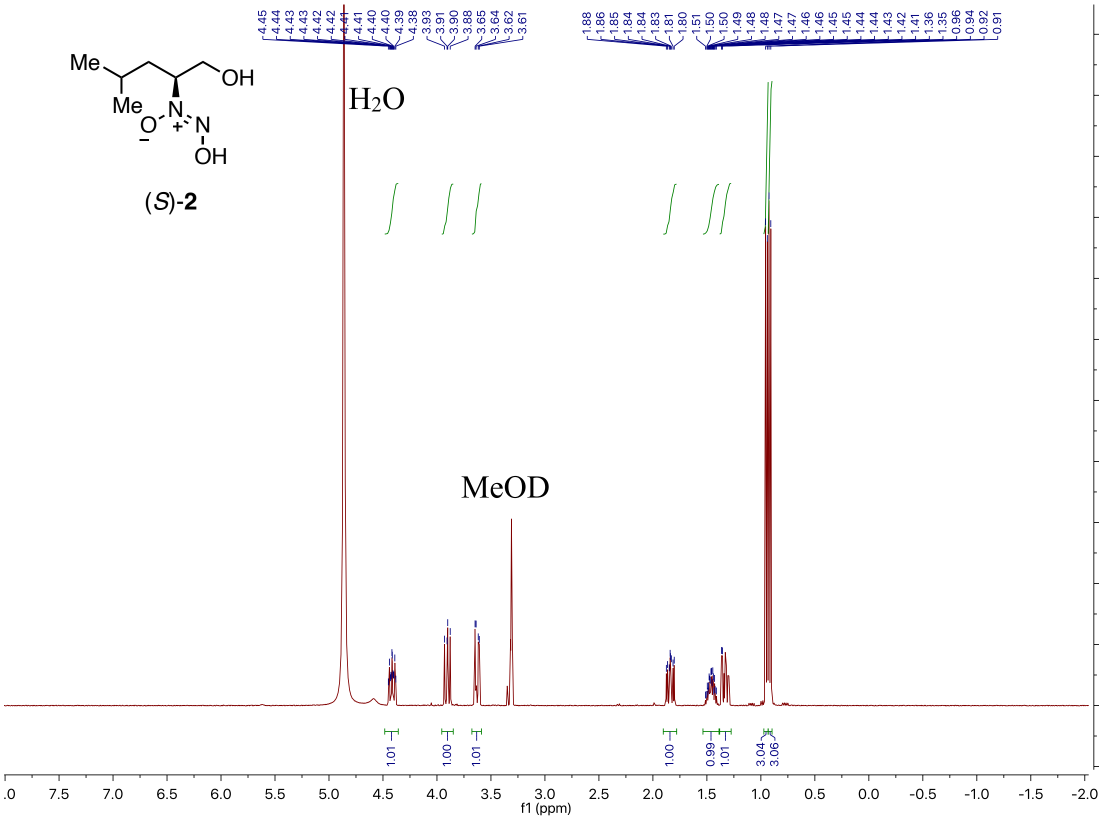


$^{13}$C-NMR Spectrum of leudiazen ((*S*)-**2**) in MeOD (400 MHz)

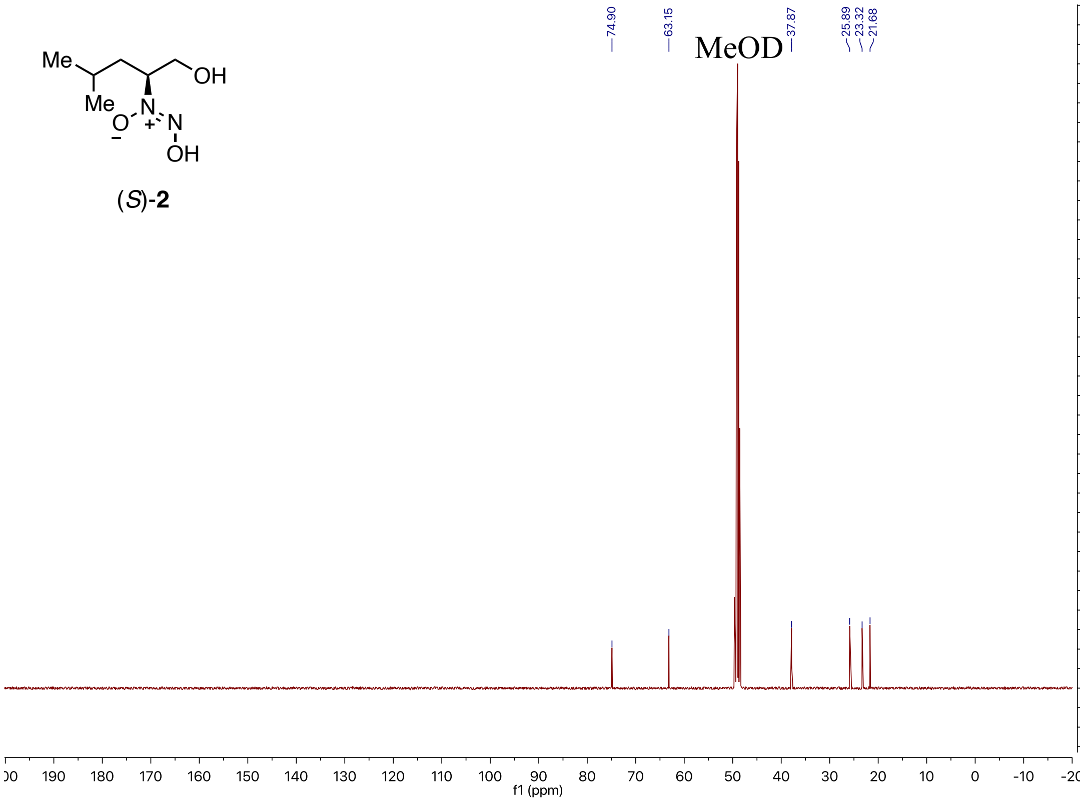

[1]H-NMR Spectrum of leudiazen ((*R*)-**2**) in MeOD (400 MHz)

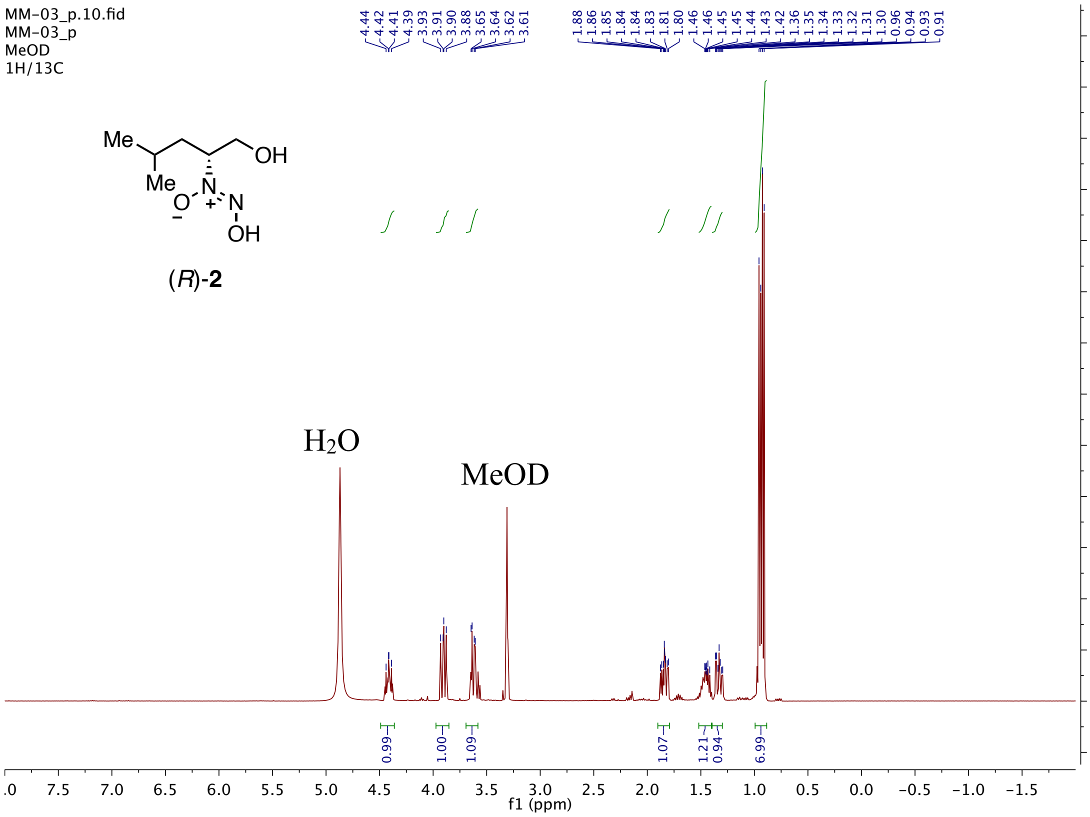


[13]C-NMR Spectrum of leudiazen ((*R*)-**2**) in MeOD (400 MHz)

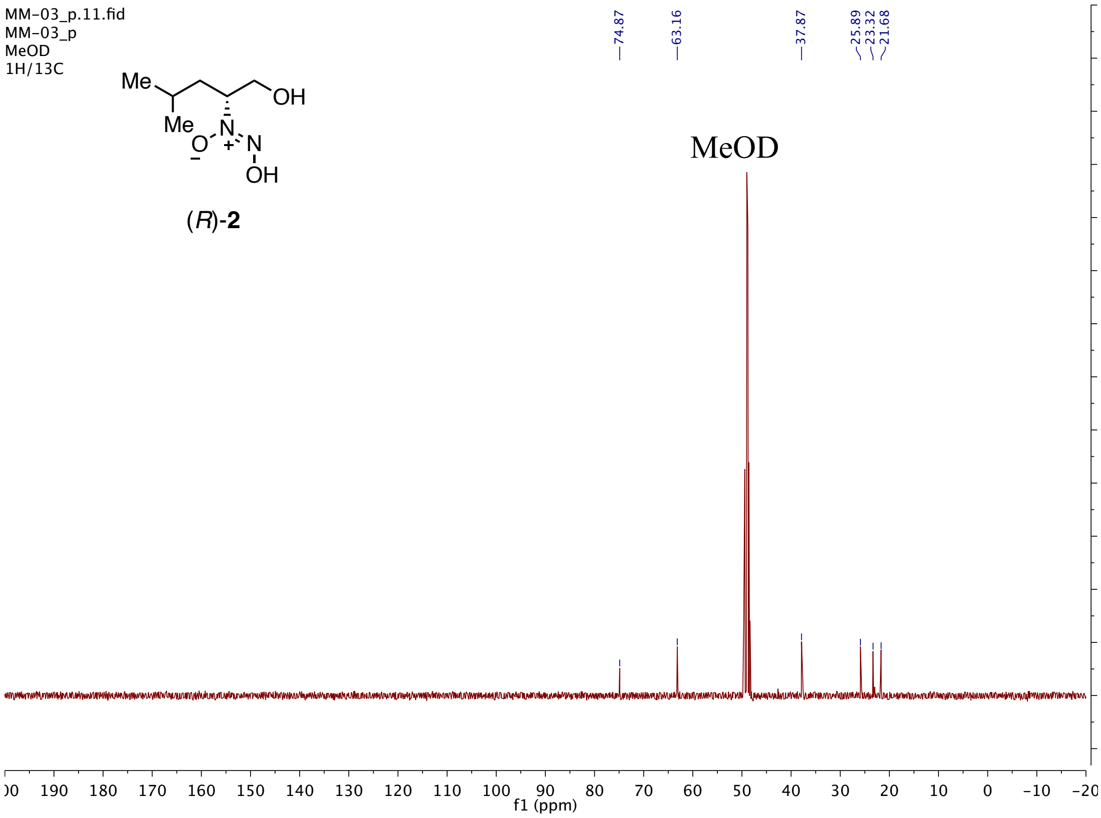

[1]H-NMR Spectrum of leudiazen (**2**) in MeOD (400 MHz)

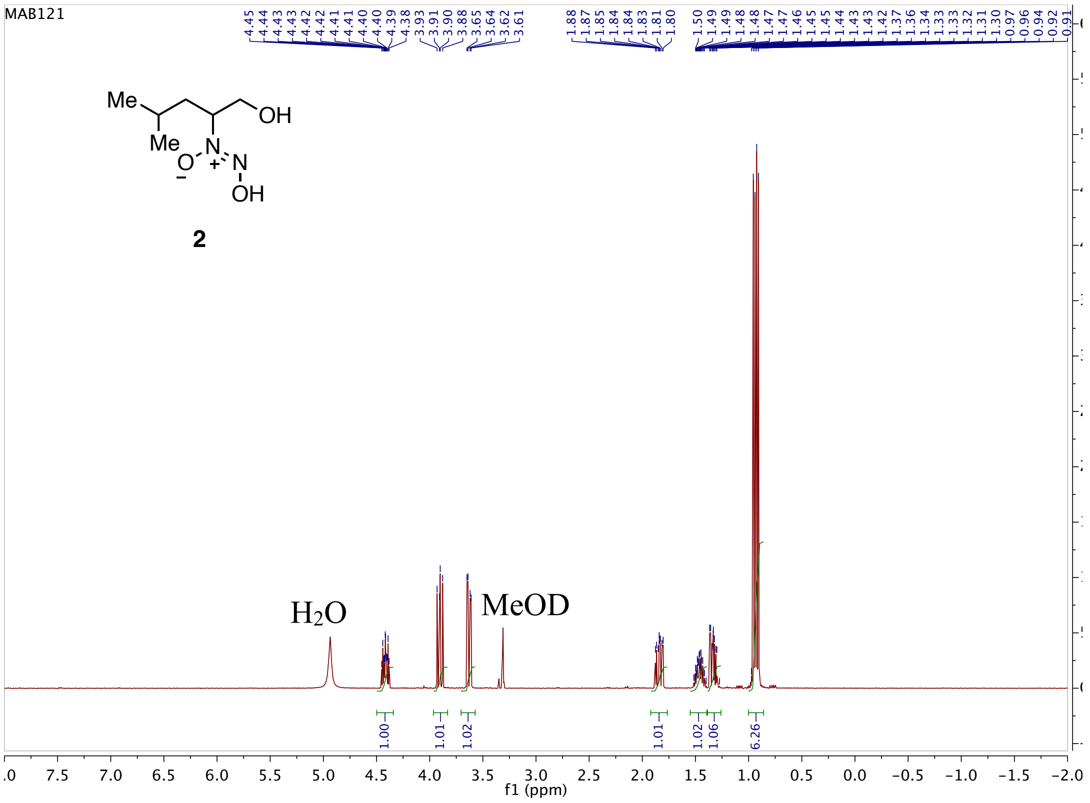


[13]C-NMR Spectrum of leudiazen (**2**) in MeOD (400 MHz)

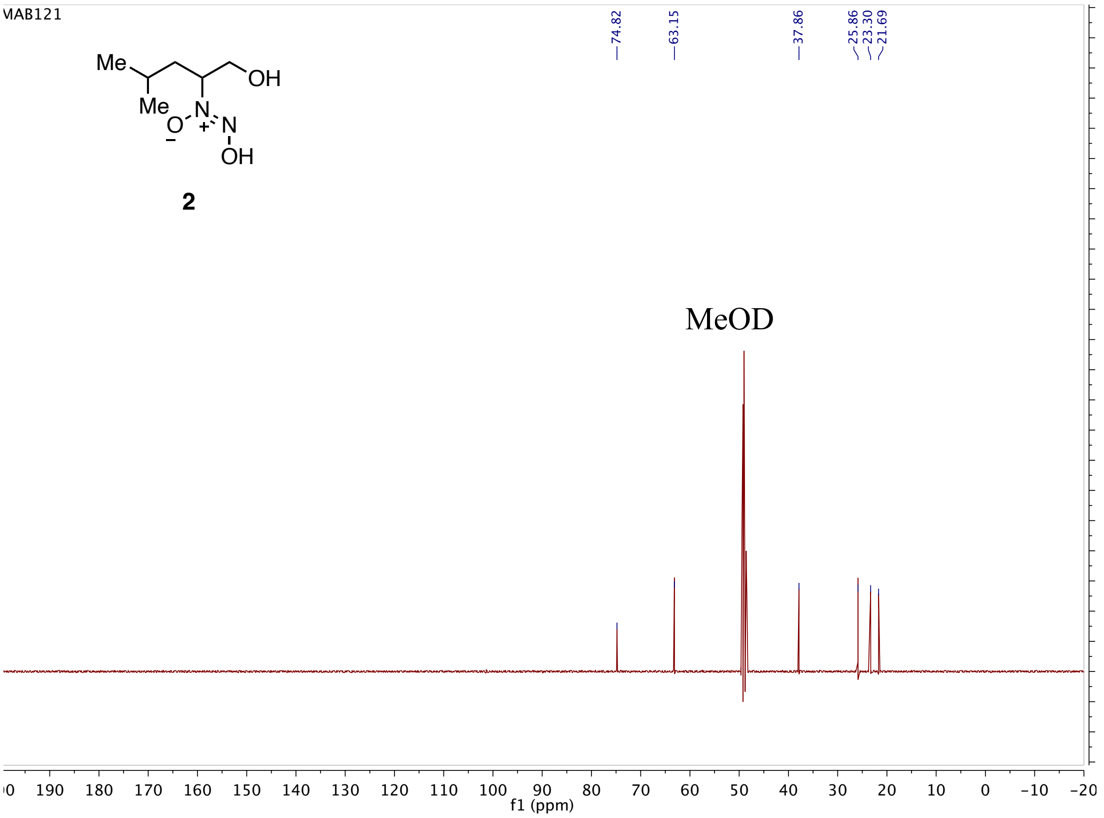

[1]H-NMR Spectrum of glydiazen (**4**) in MeOD (400 MHz)

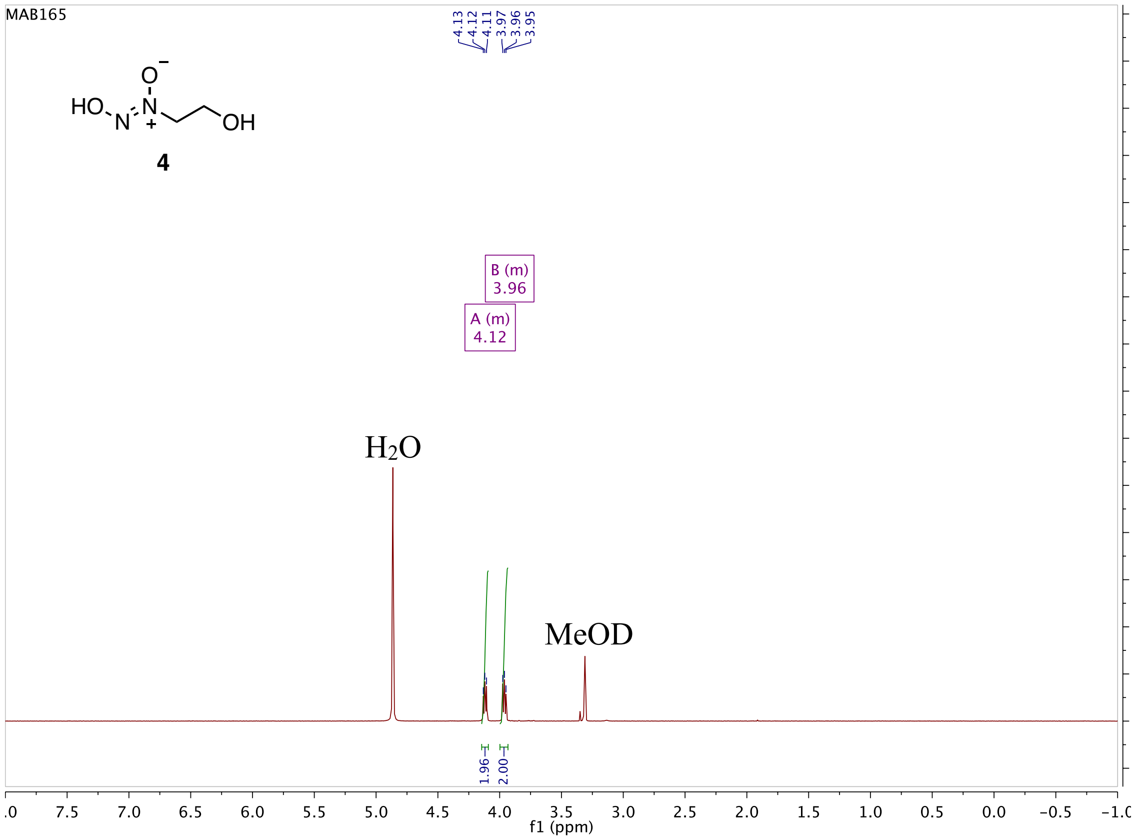


[13]C-NMR Spectrum of glydiazen (**4**) in MeOD (400 MHz)

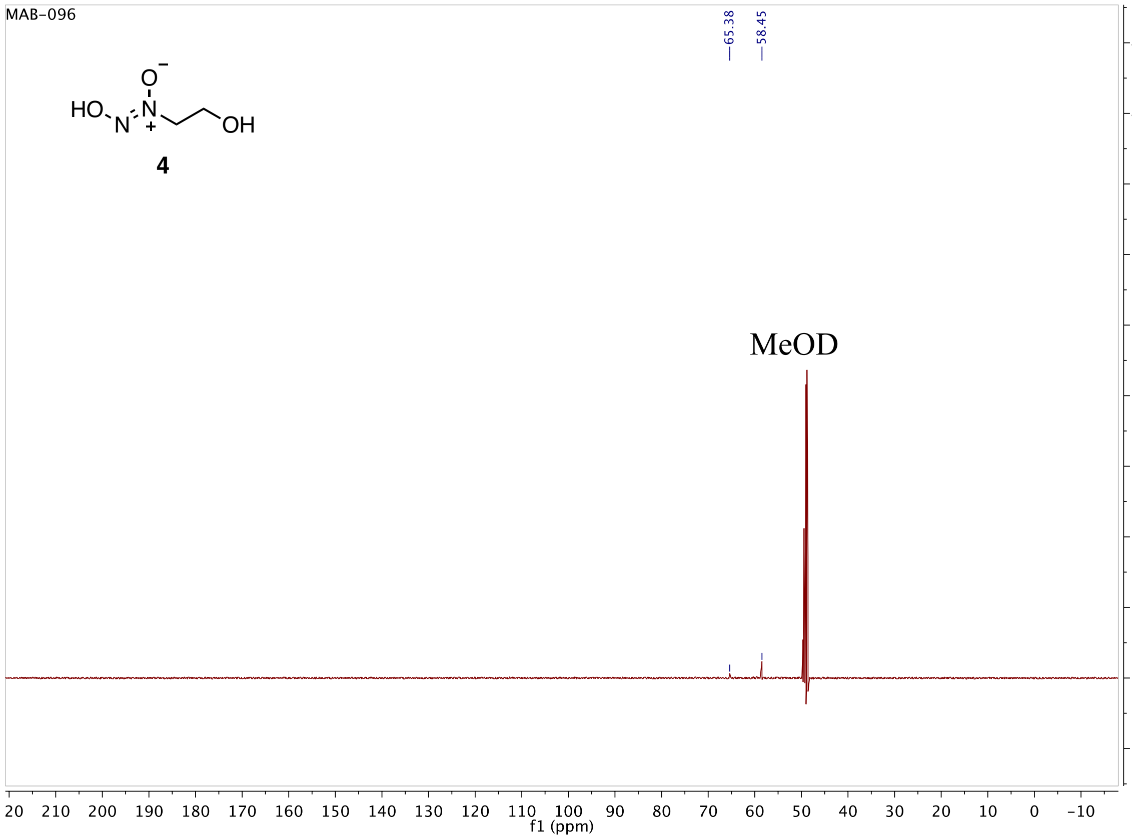

[1]H-NMR Spectrum of aladiazen (**5**) in MeOD (400 MHz)

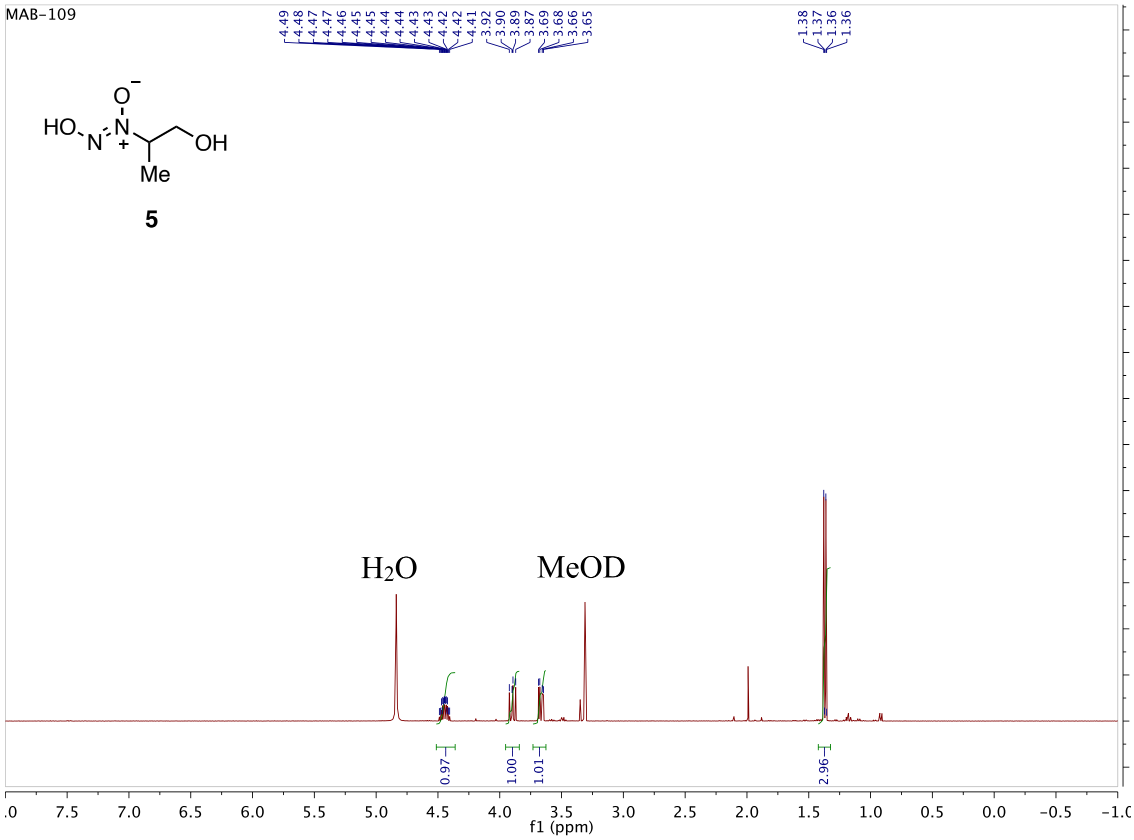


[13]C-NMR Spectrum of aladiazen (**5**) in MeOD (400 MHz)

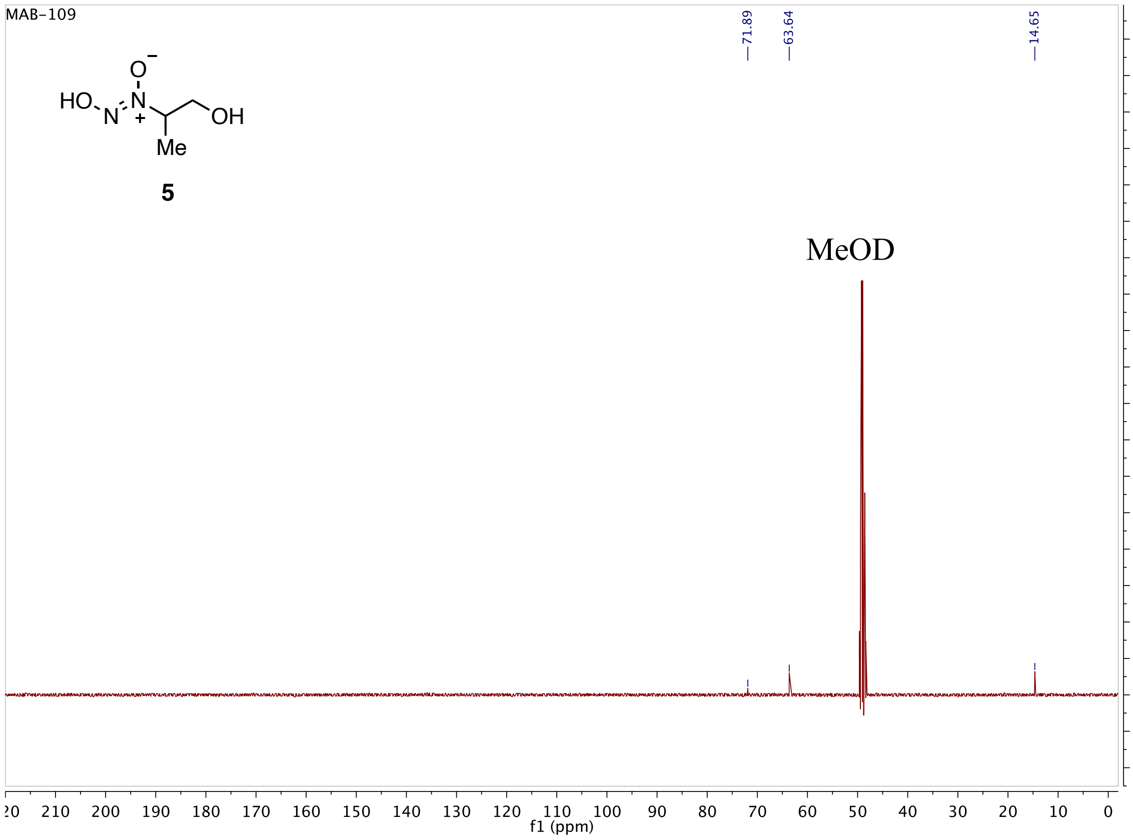

[1]H-NMR Spectrum of isoleudiazen (**3**) in MeOD (400 MHz)

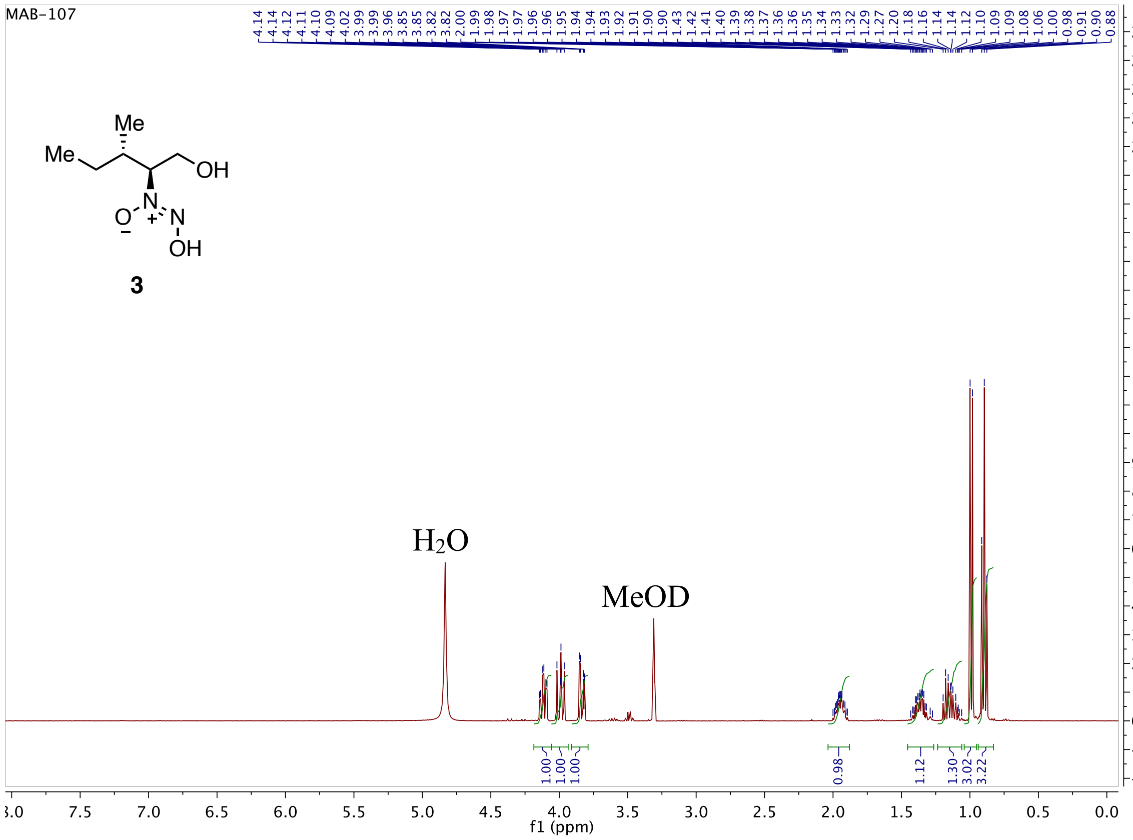


[13]C-NMR Spectrum of isoleudiazen (**3**) in MeOD (400 MHz)

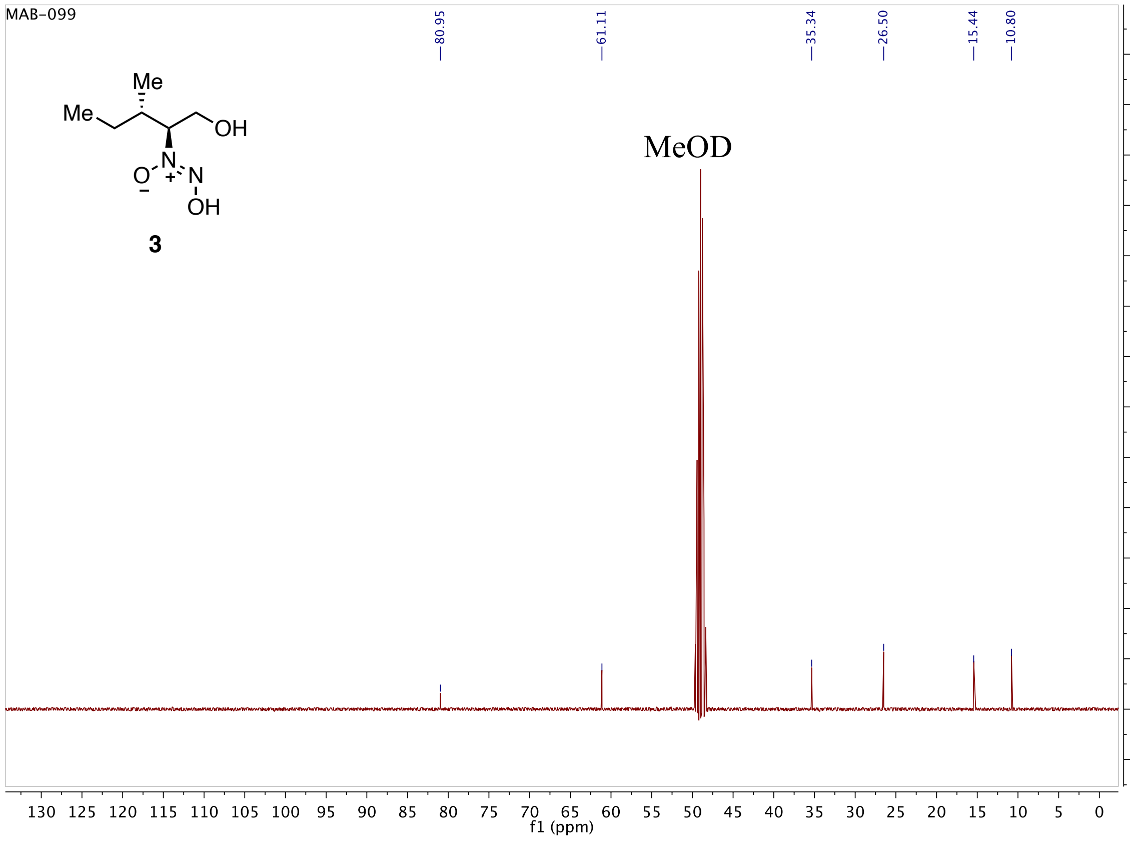

### HPLC Spectra

HPLC-UV chromatogram (254 nm) of the wild type extract, pre-purified by SPE. The separation was performed on an Agilent instrument (1100 serie) using a mobile phase composed of $H_2O$ (0.1% $HCO_2H$) and MeCN (0.1% $HCO_2H$). Four fractions were collected (from 0 to 8 min, from 8 to 9.6 min, from 9.6 to 10.5 min and from 10.5 to 25 min).

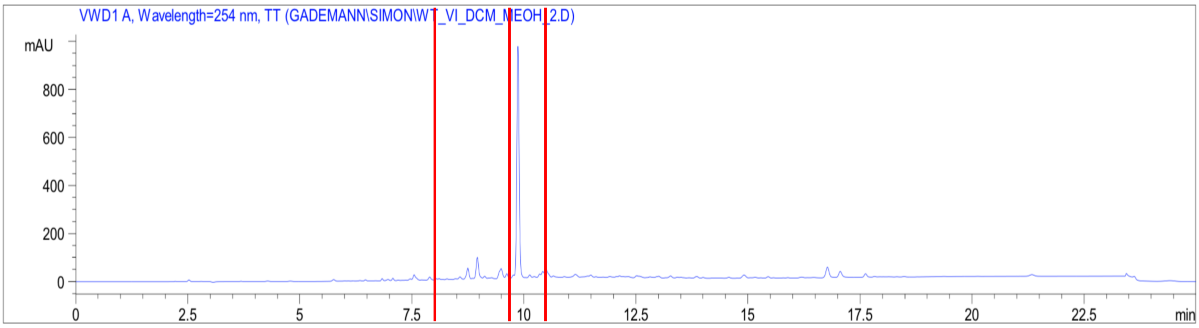


HPLC-UV chromatogram (254 nm) of the active fraction collected at from 8 to 9.6 min. The separation was performed on an Agilent instrument (1100 serie) using a mobile phase composed of $H_2O$ (0.1% $NH_4OAc$) and MeCN:$H_2O$ (95:5, 0.1% $NH_4OAc$). Four fractions were collected (from 0 to 8 min, from 8 to 9 min, from 9 to 10 min and from 10 to 25 min).

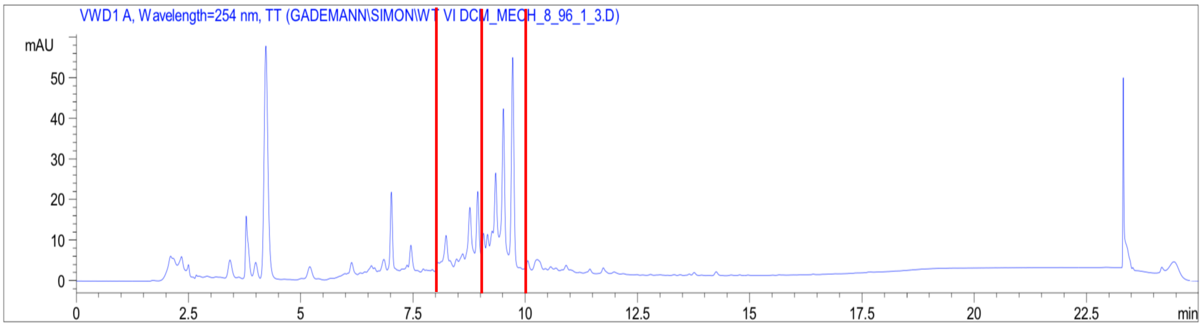

UHPLC-MS chromatograms (100-600 Da) comparison from top to bottom of the Pss extract (3 mg/mL), *Δmgo* mutant extract (3 mg/mL), and leudiazen (50 µg/mL).

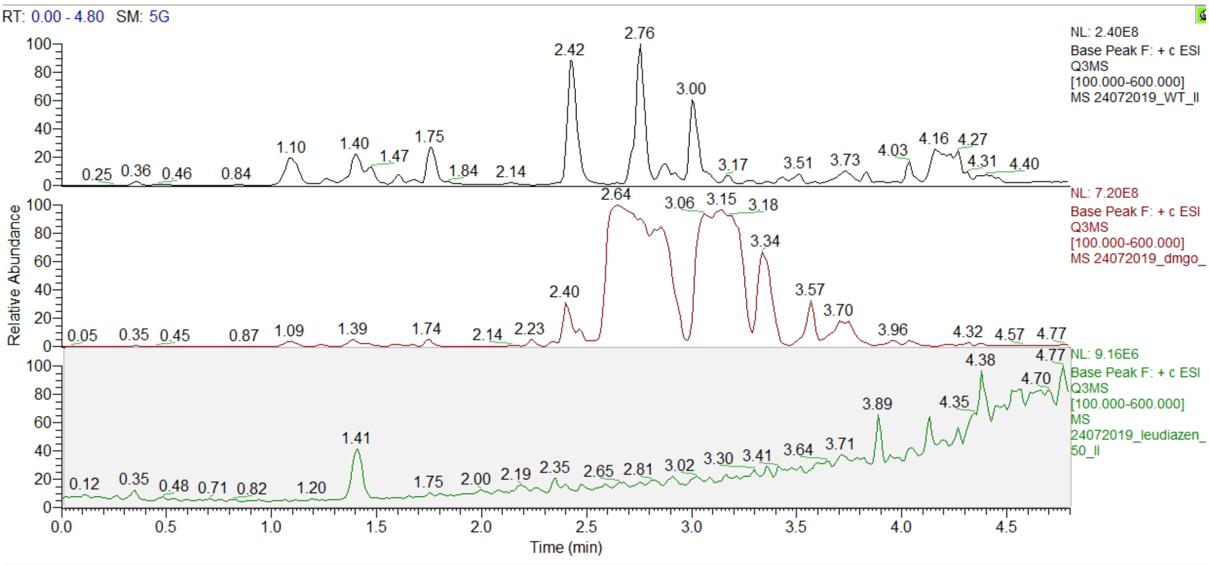
